# A Cross-Species Enhancer-AAV Toolkit for Cell Type-Specific Targeting Across the Basal Ganglia

**DOI:** 10.64898/2026.02.23.706695

**Authors:** Morgan E. Wirthlin, Avery C. Hunker, Saroja Somasundaram, M. Nathaly Lerma, William D. Laird, Victoria Omstead, Naz Taskin, Niklas Kempynck, Matthew T. Schmitz, Yuan Gao, Emma Thomas, Marcus Hooper, Yoav Ben-Simon, Refugio A. Martinez, Ximena Opitz-Araya, John Mich, Aaron Oster, Deepanjali Dwivedi, Erin Groce, Jada Roth, Bargavi Thyagarajan, Sharon Way, Avalon Amaya, Angela Ayala, Stuard Barta, Darren Bertagnolli, Madeline Bixby, Trangthanh Cardenas, Tamara Casper, Michael Clark, Nicholas Donadio, Nadezhda I. Dotson, Tom Egdorf, Erica L. Peterson, Jessica Gloe, Jeff Goldy, Conor Grasso, Warren Han, Samantha D. Hastings, Madeleine Hewitt, Daniel Hirschstein, Windy Ho, Alvin Huang, Tye Johnson, Danielle Jones, Atlas Jordan, Zoe C. Juneau, Matthew Jungert, Madhav Kannan, Jaimie Kenney, Shannon Khem, Rana Kutsal, Samantha Lee, Henry Loeffler, Nicholas Lusk, Rachel McCue, Robyn Naidoo, Dakota Newman, Nhan-Kiet Ngo, Katrina Nguyen, Sven Otto, Ben Ouellette, Alana Oyama, Nick Pena, Elliot Phillips, Christina Alice Pom, Lydia Potekhina, Shea Ransford, Christine Rimorin, Dana Rocha, Augustin Ruiz, Raymond Sanchez, Stephanie Seeman, Lucas Suarez, Mike J. Taormina, Michael Tieu, Alex Tran, Tulip E. O’Neill, Natalie Weed, Josh Wilkes, Jeanelle Ariza, Anish Bhaswanth Chakka, Forrest Collman, Nick Dee, Tyler Mollenkopf, Nadiya V. Shapovalova, Kimberly Smith, Jack Waters, Ali Williford, Shenqin Yao, Ina Ersing, Marcella Patrick, Nelson J. Johansen, Jeremy Miller, Hongkui Zeng, Ed S. Lein, Yoshiko Kojima, Gregory D. Horwitz, Boaz P. Levi, Tanya L. Daigle, Bosiljka Tasic, Trygve E. Bakken, Jonathan T. Ting

**Author notes:** These authors contributed equally (Morgan E. Wirthlin, Avery C. Hunker).

## Abstract

The mammalian basal ganglia (BG) orchestrate motor, cognitive, and affective functions, yet cell type-specific genetic access remains limited, especially beyond rodents. Key structures implicated in movement and psychiatric disorders, including pallidum, subthalamic nucleus, and dopaminergic midbrain, lack scalable tools for cross-species targeting. Here, we present a comprehensive enhancer-AAV library enabling selective labeling and manipulation of major BG neuronal populations: striatal projection neuron subtypes, pallidal and subthalamic neurons, and midbrain dopaminergic and GABAergic populations. Using an evolutionarily informed discovery pipeline, we identified enhancers targeting canonical, non-canonical, and disease-relevant cell types, with validation demonstrating robust cross-species conservation of specificity between mouse and macaque. Computational modeling revealed sequence features predictive of in vivo performance, including motif grammar, chromatin accessibility, and evolutionary conservation, and identified distinct regulatory architectures across glial, projection, and interneuron lineages. This work establishes a comprehensive cross-species viral toolkit for the BG, unlocking previously inaccessible cell types for circuit dissection.

## INTRODUCTION

The basal ganglia (BG) form a set of interconnected subcortical nuclei that collectively orchestrate motor execution, reinforcement learning, habit formation, and affective control (Graybiel et al. 1994; Kreitzer and Malenka 2008). The BG circuitry responsible for these processes involves multiple hierarchically organized loops linking the cortex, thalamus, and brainstem and involving excitatory, inhibitory, and dopaminergic signaling pathways (Alexander et al. 1986; Bolam et al. 2000). Dysfunction within these networks underlies a broad range of neurological and psychiatric disorders, including Parkinson’s disease, dystonia, Huntington’s disease, Tourette syndrome, obsessive-compulsive disorder, and addiction (Graybiel and Rauch 2000; Haber and Rauch 2010; Obeso et al. 2008; Albin et al. 1989).

Over the past two decades, cell type-specific transgenic mouse lines have revolutionized our understanding of striatal circuits. Cre-driver and reporter lines from the Gene Expression and Nervous System Atlas (GENSAT) project (Gong et al. 2003) and related projects have enabled genetic access to the canonical direct (D1) and indirect (D2) pathway medium spiny neurons (Gerfen et al. 2013; Bateup et al. 2010) among many other cell types, enabling extensive interrogation of classical BG circuit models. However, the reach of these resources is limited: they are often transgenic lines restricted to mouse studies, rely on complex breeding or intersectional labeling strategies (Zeng and Sanes 2017), or fail to fully capture the molecular and anatomical diversity of BG cell types. Transgenic approaches are particularly challenging for larger mammals such as NHPs (Jennings et al. 2016), which remain essential for modeling human motor and cognitive functions. As a result, major nodes of the primate BG—including the globus pallidus, subthalamic nucleus (STN), and midbrain neuron populations—have remained inaccessible to precise genetic perturbation as compared to the plethora of available genetic tools and targeting strategies for BG cell types in the mouse model (Gerfen et al. 2013; Mastro et al. 2014; Lammel et al. 2015; Parolari et al. 2021).

Adeno-associated virus (AAV) vectors that employ cell type-selective enhancers to drive the precise delivery of their cargo offer a promising avenue to overcome these limitations (Chan et al. 2017; Graybuck et al. 2021; Hrvatin et al. 2019). Enhancer-AAVs harness regulatory DNA sequences that direct transcription within specific neuronal subclasses, enabling targeted expression without reliance on germline engineering in transgenic mice. When combined with large-scale, single-cell transcriptomic and epigenomic datasets, enhancer discovery can be systematically guided by cell type-specific chromatin accessibility (Graybuck et al. 2021; Lawler et al. 2022; Hunker et al. 2025; Kussick et al. 2025; Ben-Simon et al. 2025). This strategy has already yielded highly specific tools for the striatum, where validated enhancer-AAVs now rival or surpass traditional transgenic lines in precision and flexibility (Hunker et al.2025). Yet the BG as a whole encompass a far wider array of neuronal types, including inhibitory and excitatory projection neurons, diverse interneuron subclasses, and dopaminergic and GABAergic midbrain populations (Johansen et al. 2025; He et al. 2021). These types differ profoundly in developmental origin, connectivity, and transcriptional identity, and contribute critically to the functional processes essential for understanding BG function in normal and disease states (Crittenden and Graybiel 2011). Extending enhancer-based targeting across this full ensemble represents a central unmet need for both basic and translational research.

To address this challenge, we developed the Cross-species Enhancer Ranking Pipeline (CERP), a cross-species enhancer discovery and validation pipeline for the BG that integrates single-nucleus RNA-seq and ATAC-seq data from multiple mammalian species with predictive sequence modeling and large-scale *in vivo* screening. Using CERP, we identified regulatory elements active in 21 cell populations spanning the striatum, pallidum, subthalamic nucleus, and dopaminergic midbrain (**Fig. 1**, **Table S1**). Candidate enhancers were drawn from mouse, macaque, and human open chromatin datasets, prioritized by evolutionary conservation, motif composition, and transcription factor (TF) occupancy, which were then cloned into AAV vectors for systematic testing *in vivo*. The resulting enhancer-AAV library enables selective labeling and manipulation of canonical medium spiny neuron (MSN) subclasses, as well as pallidal projection neurons, subthalamic nucleus glutamatergic neuron types, and dopaminergic and GABAergic neurons of the substantia nigra and ventral tegmental area. Importantly, these tools achieve high specificity in both rodents and NHPs, establishing the first comprehensive cross-species genetic toolkit for dissecting BG circuits.

**Figure 1:**
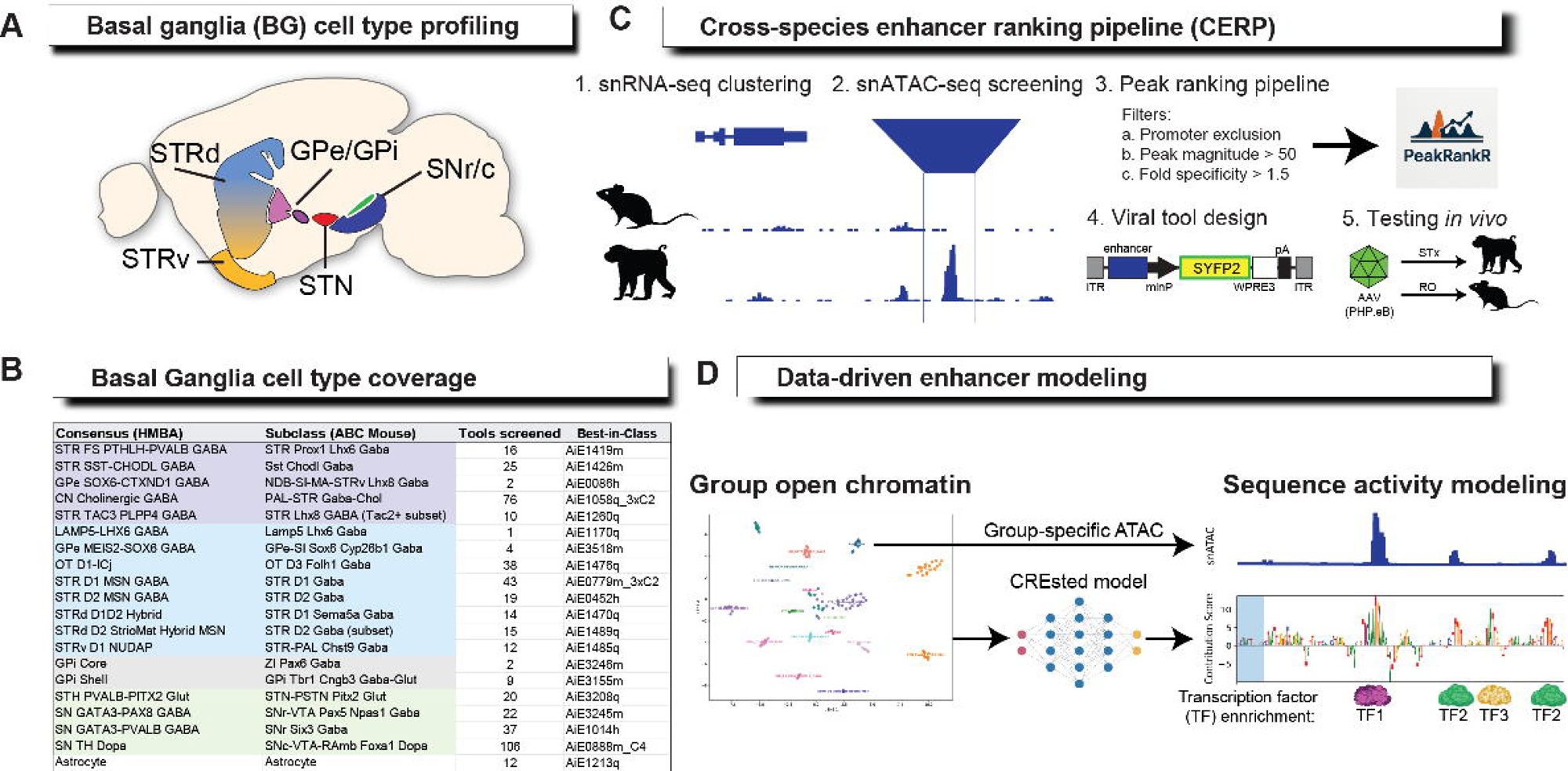
Overview of the basal ganglia enhancer-AAV toolkit and Cross-species Enhancer Ranking Pipeline (CERP). (**A**) Anatomical scope of the study, encompassing the major BG nuclei targeted in this work: dorsal and ventral striatum (STRd, STRv), globus pallidus externa and interna (GPe, GPi), subthalamic nucleus (STN), and substantia nigra/ventral tegmental area (SN/VTA). (**B**) Summary of BG cell type coverage achieved in this study. Targets are anchored to Human and Mammalian Brain Atlas (HMBA) consensus BG taxonomy nomenclature and mapped explicitly to mouse class and subclass labels from the ABC mouse brain atlas (see Supplementary Table 1). (**C**) The Cross-species Enhancer Ranking Pipeline (CERP) uses cell type-resolved snRNA-seq and snATAC-seq datasets from mouse and macaque to identify regulatory sequences for use in viral tools. Differentially accessible chromatin peaks are ranked using a feature-based prioritization strategy emphasizing cell type specificity while excluding promoter-proximal and low-strength candidates. Selected enhancers are cloned upstream of a minimal promoter in a standardized AAV backbone and tested *in vivo* using high-throughput retroorbital (RO) delivery in mouse and stereotaxic (STX) delivery for spatially restricted targeting, including in NHP. (**D**) Data-driven modeling framework used to interpret validated enhancers and identify sequence features predictive of tool performance. Group-resolved chromatin accessibility is integrated with sequence-based activity modeling to evaluate transcription factor motif composition, predicted regulatory contributions, and evolutionary constraint, enabling mechanistic inference of *cis*-regulatory logic underlying cell type-specific enhancer activity and informing cross-species comparisons.

By coupling *in vivo* functional validation with computational modeling, we identify DNA sequence features predictive of enhancer performance—including motif combinations, local chromatin accessibility, and evolutionary constraint—that together define cell type-specific regulatory codes. Glial, interneuron, and projection neuron lineages each exhibit distinct enhancer architectures and TF grammars (Johansen et al. 2025), illuminating the molecular principles that partition cell type specificity across BG circuits. Comparative analysis of orthologous sequences further reveals how conserved and divergent enhancer activity contributes to the regulatory evolution of BG cell type identity, bridging mechanistic genomics with functional neuroanatomy.

Collectively, these findings establish a unified, evolutionarily informed approach to design, validate, and interpret enhancer-based viral tools across species. The BG Enhancer-AAV Toolkit extends cell type-specific access from the striatum to all major nuclei of the BG, enabling precise experimental dissection of circuits implicated in movement, motivation, and neuropsychiatric disease. By coupling cross-species enhancer discovery with predictive modeling, this work provides both a practical and conceptual foundation for future efforts to engineer regulatory-element-driven gene therapies and to chart the genomic underpinnings of brain evolution.

## RESULTS

### Cross-species enhancer discovery and atlas-aligned targeting across the BG

The basal ganglia encompass the dorsal and ventral striatum (STRd, STRv), globus pallidus externa and interna (GPe/GPi), subthalamic nucleus (STN), and substantia nigra / ventral tegmental area (SN/VTA) (**Fig. 1A**). Because BG cell types are labeled inconsistently in the literature and across species-specific atlases, we employed the HMBA consensus nomenclature (Johansen et al. 2025) and provided explicit mappings to mouse class and subclass labels used in the ABC whole mouse brain atlas (Yao et al. 2023) (**Fig. 1B**). The 21 atlas-defined targeted populations span glial, inhibitory, excitatory, and neuromodulatory lineages and cover both canonical BG types and rare or anatomically restricted types that have been difficult to access with existing genetic resources. Within striatum, targets included the canonical STR D1 and STR D2 medium spiny neurons (MSNs) and atlas-resolved MSN-adjacent or hybrid populations, including STRd D1D2 Hybrid and STRv D1 NUDAP populations (He et al. 2021), known collectively in the literature as eccentric MSNs (eMSNs) (Saunders et al. 2018; Gayden et al.2023), OT D1-ICj granule cells, as well as the recently defined STRd D2 StrioMat Hybrid MSNs (Johansen et al. 2025). We also focused on major STR interneuron subclasses including STR FS PTHLH-PVALB GABA and STR SST-CHODL GABA.

Beyond striatum, we targeted major GABAergic projection neuron classes of pallidum, including GPe MEIS2-SOX6 GABA and GPe SOX6-CTXND1 GABA, as well as BG output populations in GPi (GPi Core and GPi Shell) and the glutamatergic STN lineage (STH PVALB-PITX2 Glut). Finally, we included midbrain GABAergic and dopaminergic neuron populations that gate BG output and provide neuromodulatory control, including anterior and posterior SN GABA groups (SN GATA3-PVALB GABA and SN GATA3-PAX8 GABA, respectively) and multiple SN TH dopaminergic targets, encompassing pan-dopaminergic populations and dorsal tier and ventral tier-biased subtypes aligned to spatial and molecular subdivisions (Gernert and Löscher 2001; Fiorenzano et al. 2021). Together, these targets define a BG-wide testbed for enhancer discovery that spans multiple developmental origins and neurotransmitter classes while remaining grounded in a unified cross-species cell type framework.

To generate tools across this breadth of nuclei and cell types, CERP was applied to a BG cell type taxonomy that was defined in a companion study (Johansen et al. 2025) based on single nucleus multiomic data. CERP identified cell type-enriched open chromatin peaks and ranked peaks using a feature-based prioritization strategy that emphasizes cell type-resolved accessibility, open chromatin strength, and cross-species sequence conservation, while excluding promoter-proximal elements and low-specificity candidates (**Fig. 1C** and **Methods**). Selected peaks were cloned upstream of a minimal promoter in a standardized AAV backbone, packaged into viral particles and tested by mouse *in vivo* systemic administration (retro-orbital delivery) for high-throughput screening and stereotaxic injection for spatially restricted delivery, including in NHP. This design enables systematic comparison of enhancer performance across cell type targets under controlled conditions, while providing a scalable path from chromatin accessibility to validated viral reagents. We experimentally evaluated 514 candidate enhancers targeting all BG areas. This systematic screen was undertaken to identify, for each target population, a set of best-in-class enhancers, which we defined as those exhibiting highest on-target specificity with minimal off-target labeling within our experimental framework, and that together constitute a ready-to-use toolkit for precise experimental access to BG cell types.

Finally, we implemented a data-driven modeling strategy to interpret validated enhancers and to identify sequence-level features that predict tool performance (**Fig. 1D**). We integrated cell type-resolved chromatin accessibility with sequence-based activity modeling to assess open chromatin features, combinations of transcription factor binding motifs, and evolutionary constraints that distinguish on-target from off-target enhancer-AAV candidates within the validated library. This approach bridges enhancer discovery and validation to mechanistic inference of *cis*-regulatory logic across species and quantifies to what degree enhancer function depends on overall sequence identity versus conserved regulatory grammar.

Together, this resource defines a comprehensive BG resource that integrates cross-species enhancer discovery, atlas-aligned cell type targeting, and predictive regulatory modeling, establishing both a scalable toolkit for circuit access and a principled foundation for interpreting the *cis*-regulatory logic underlying BG cell type specificity.

### Large-scale *in vivo* screening identifies high-specificity enhancer-AAVs across BG cell types

To evaluate the ability of cell type-specific enhancers to drive targeted expression across the BG, each candidate enhancer sequence from our CERP-BG library was cloned into an SYFP2 reporter vector, packaged into the blood brain barrier-penetrant AAV with PHP.eB capsid (Chan et al. 2017), and delivered to mice by retro-orbital (RO) injection. We then used a one at a time enhancer-AAV screening pipeline, divided into primary and secondary evaluation stages, to narrow down the top candidates across BG regions (**Fig. 2A**), similar to as described previously (Hunker et al. 2025).

**Figure 2:**
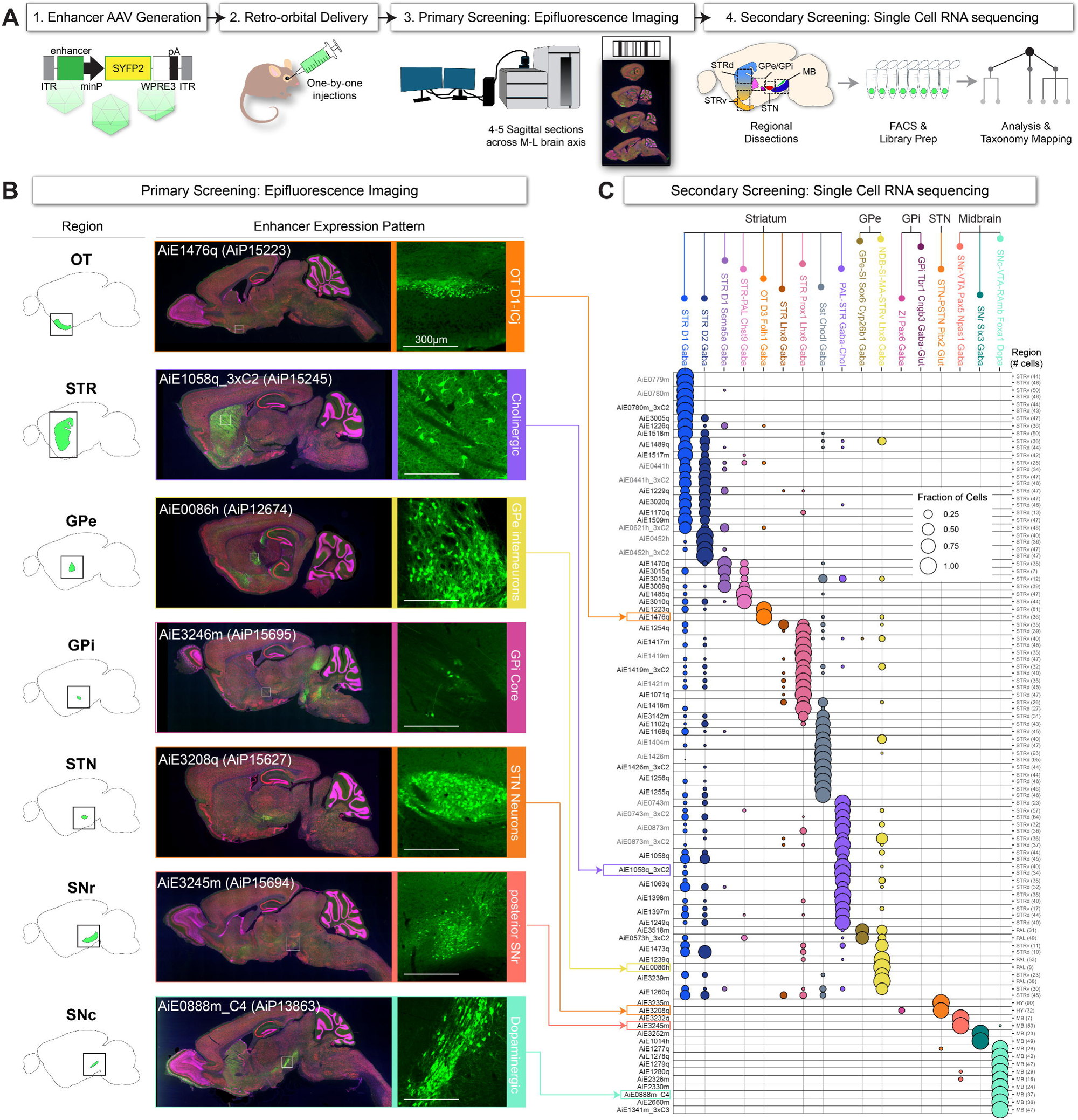
Large-scale *in vivo* screening of BG enhancer-AAVs in mouse. (**A**) Schematic of the *in vivo* enhancer screening pipeline. Candidate enhancers are cloned into AAV vectors driving SYFP2, delivered retro-orbitally (RO) in mouse, and assessed by primary epifluorescence imaging of sagittal brain sections. Labeled cells from dissected BG regions are subsequently isolated for secondary screening by single-cell RNA sequencing (SSv4) and mapped to HMBA BG consensus atlas-defined cell types. (**B**) Representative epifluorescence images from primary screening illustrating enhancer-driven SYFP2 expression across BG regions. For each example, left panels show anatomical regions of interest in sagittal mouse brain atlas sections and right panels show full sagittal SYFP2 sections with higher-magnification insets of the indicated region. Regions shown include olfactory tubercle (OT), striatum (STR), globus pallidus externa (GPe) and interna (GPi), subthalamic nucleus (STN), substantia nigra pars reticulata (SNr), and dopaminergic midbrain, substantia nigra pars compacta (SNc). Scale bars, 300 μm. (**C**) Secondary screening by SSv4. Bubble plot showing the distribution of SYFP2-positive cells across atlas-defined BG cell types for each tested enhancer. Dot size indicates the fraction of labeled cells assigned to each cell type, and colors denote cell type identity grouped by anatomical region: dorsal and ventral striatum (STRd, STRv), pallium (PAL), hypothalamus (HY), and midbrain (MB). The number of sequenced cells per region is indicated at right.

We used primary screening as a first-pass assessment of enhancer-driven reporter expression in the mouse brain. Sagittal sections from enhancer AAV-injected mouse brains were imaged for SYFP2 native fluorescence at 4-5 different medial-lateral depths (**Fig. 2B**). To initially evaluate on-target expression, we used different cell features depending on cell type target. For example, striatal cholinergic-targeted enhancers were identified by the large soma size of striatal cholinergic neurons, whereas STN enhancers could be clearly identified by regional SYFP2 expression. For various BG cell types, we also utilized the Allen Institute mouse spatial ABC Atlas as a guide for determining cell type spatial patterns (Yao et al.2023).

Following primary screening, we selected 36 of our most promising candidates across all BG regions for evaluation in our secondary screening pipeline (**Fig. 2C, Table S1**), where we quantified cell type specificity using single cell RNA-sequencing (scRNA-seq). We isolated and sorted whole cells from each enhancer AAV-injected mouse brain for SYFP2 fluorescence and performed scRNA-seq by SMART-Seq v4 (SSv4) (**Methods**). To determine cell type identity, we bioinformatically mapped sequenced cells to the ABC atlas whole mouse brain taxonomy at the subclass level using Map-My-Cells algorithm (**Methods**) (Yao et al. 2023). To round out our BG enhancer collection we revisited 15 of our previously published best-in-class dorsal striatum enhancers (Hunker et al. 2025) and extended our analysis to include ventral striatum validation. Across regions,we demonstrated that best-in-class enhancers drove expression with >70% on-target specificity, with exemplary enhancers achieving >90% on-target specificity (**Fig. 2C**).

These data extend the reach of prior striatal enhancer resources to encompass the full diversity of MSN and MSN-adjacent types formalized in the HMBA taxonomy (Johansen et al. 2025).

Beyond striatal populations, CERP-derived enhancers enabled, for the first time, systematic genetic access to the primary cell types of the small nuclei of the BG—structures for which no enhancer-based viral tools have previously existed. In the GPe, electrophysiological and molecular studies over the past decade have established a fundamental dichotomy between two major GABAergic projection neuron classes: the “prototypic” neurons, which express *Nkx2-1* and *Lhx6*, fire tonically at high rates, and project to STN and downstream BG output nuclei, and the “arkypallidal” neurons, which express *FoxP2* and *Meis2*, fire at lower rates, and project exclusively back to the striatum (Mallet et al. 2012; Abdi et al.2015). In the HMBA consensus taxonomy, the prototypic population corresponds to GPe SOX6-CTXND1 GABA (mouse ABC Atlas subclass NDB-SI-MA-STRv Lhx8 Gaba), while the arkypallidal population corresponds to GPe MEIS2-SOX6 GABA (mouse ABC Atlas subclass GPe-SI Sox6 Cyp26b1 Gaba). We identified enhancers with clearly separable activity in these two classes: best-in-class prototypic GPe enhancers (AiE0086h and AiE1239q) drove reporter expression with 100% and 92% on-target specificity, respectively, while enhancer (AiE3518m) exhibited biased arkypallidal neuron labeling with 61% specificity for GPe-SI Sox6 Cyp26b1 Gaba vs. 35% specificity for NDB-SI-MA-STRv Lhx8 Gaba neurons (**Fig. 2C**). Further optimization of AiE3518m may lead to improved specificity for GPe arkypallidal neuron targeting. GPi populations (GPi Core and GPi Shell) were also targeted but were more challenging to quantify relating to the weak and sparse labeling in this small BG region. Nonetheless, primary screening data analysis supports putative on-target labeling of GPi core neurons (mouse ABC Atlas subclass ZI Pax6 Gaba) for enhancer AiE3246m (**Fig. 2B**), and putative on-target labeling of GPi shell neurons (mouse ABC Atlas subclass GPi Tbr1 Cngb3 Gaba-Glut) for enhancer AiE3155m. Further enhancer optimization for GPi subclasses remains an important goal.

In the substantia nigra pars reticulata (SNr), the principal GABAergic output nucleus of the BG, recent developmental and molecular studies have established that SNr neurons comprise at least two major subtypes of distinct embryological origin: anterior SNr neurons, which derive from *Nkx6-2*-expressing progenitors in the ventrolateral midbrain–diencephalon junction and are marked by *Six3* and *Foxp1*, and posterior SNr neurons, which originate from *Nkx6-1*-expressing progenitors and are marked by *Pax5* (Lahti et al. 2016; Partanen and Achim 2022). In the HMBA taxonomy, the anterior subtype corresponds to SN GATA3-PVALB GABA (mouse ABC subclass: SNr Six3 Gaba), while the posterior subtype corresponds to SN GATA3-PAX8 GABA (mouse ABC subclass: SNr-VTA Pax5 Npas1 Gaba). CERP-derived enhancers achieved selective labeling of these populations with mutual exclusivity: best-in-class anterior SNr enhancer (AiE3252m) drove expression at 100% on-target specificity, while posterior SNr enhancers (AiE3245m and AiP3232q) labeled this distinct population with 100% on-target specificity (**Fig. 2B,C**).

For the STN, the sole glutamatergic nucleus of the BG and a major target for deep brain stimulation in Parkinson’s disease (Okun 2012), molecular identity is defined by co-expression of *Pitx2* and *Slc17a6 (*Vglut2) (Martin et al. 2004; Wallén-Mackenzie et al. 2020). Enhancers AiE3208q and AiE3235m targeting the STH PVALB-PITX2 Glut class (mouse ABC Atlas subclass STN-PSTN Pitx2 Glut) achieved 88% and 100% on-target specificity by scRNA-seq, respectively, with near exclusive labeling of STN neurons in the whole mouse brain (**Fig. 2B,C**). Together, these results demonstrate that enhancer selection based on cell type-resolved chromatin accessibility yields reproducible, high-specificity tools across BG cell types, extending genetic access from the well-studied dorsal and ventral striatum to additional pallidal, nigral, and subthalamic populations for which viral genetic targeting tools have been lacking.

### Striatal interneuron enhancer specificity is conserved across species

We next evaluated enhancer performance in macaques *in vivo*, focusing first on striatal interneuron subclasses with robust cross-species transcriptional homology. Two groups—the STR SST-CHODL GABA interneurons (**Fig. 3A-D**) and the STR PTHLH-PVALB GABA fast-spiking interneurons (**Fig. 3E-H**)—served as stringent test cases due to their distinctive molecular signatures, sparse distribution relative to dominant striatal MSN populations, and clear roles in striatal microcircuit organization (Saunders et al. 2015; Gokce et al. 2016; Graybiel and Grafton 2015; Muñoz-Manchado et al. 2018). Stereotaxic injection of enhancer-AAVs into mouse dorsal striatum and macaque caudate revealed preservation of cell type specificity across species (**Fig. 3A–B, E–F**).

**Figure 3:**
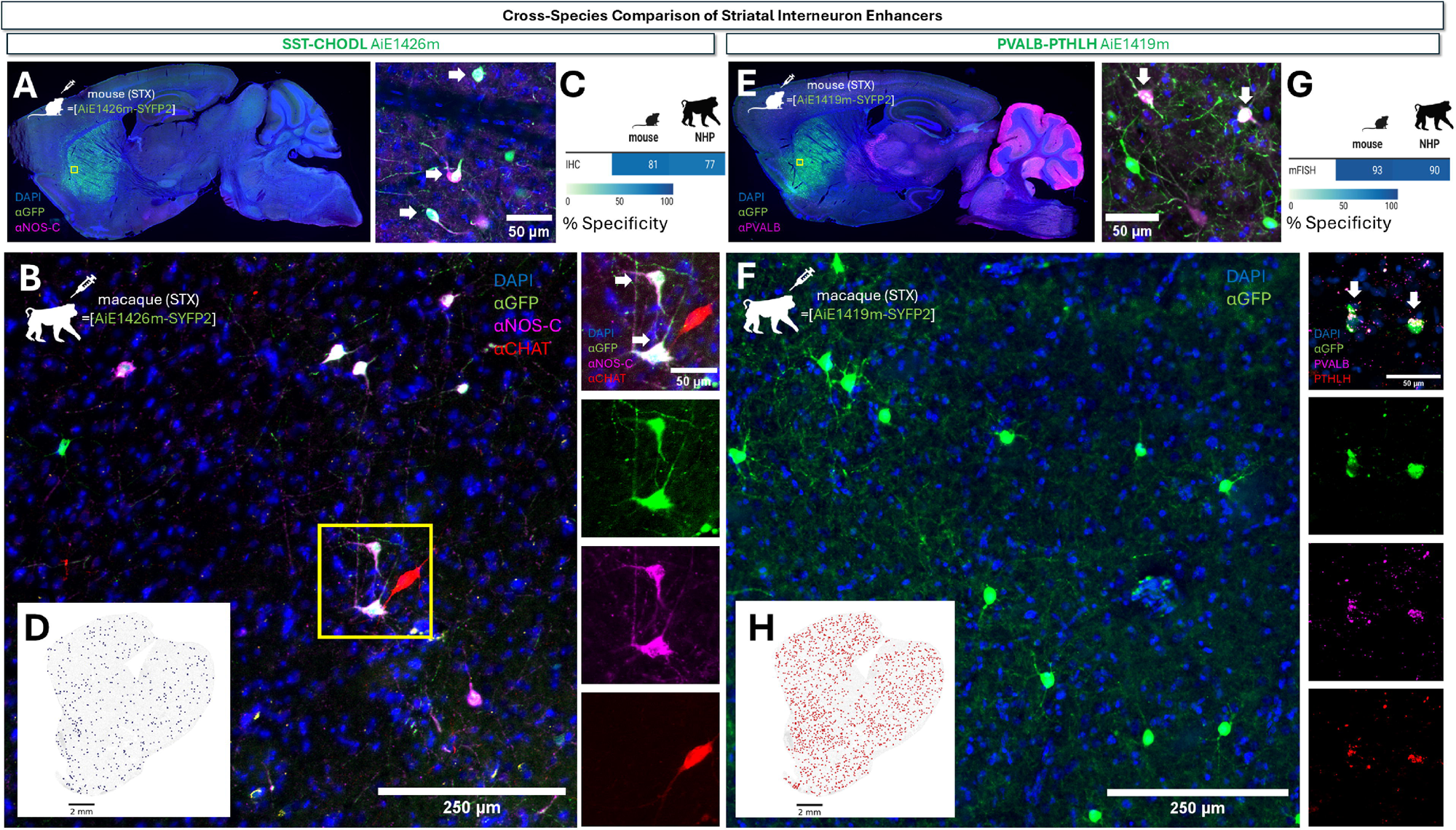
Cross-species comparison of striatal interneuron populations in mouse and macaque striatum. (**A-B**) Stereotaxic injection of AAV vector AiP15050 (best in class SST-CHODL enhancer AiE1426m driving SYFP2) in mouse dorsal Str and macaque Cd. The total viral dose was 5E+8vg for mouse dorsal Str and 1E+11vg for macaque caudate. (**A**) Sagittal section of mouse brain showing DAPI, αGFP, and αNos-C staining. Inset represents the yellow boxed region as shown. (**B**) Coronal section of macaque Cd region showing DAPI, αGFP, αNos-C, αCHAT staining. The image panels at right show magnified views for composite and individual channels for αGFP (green), αNos-C (magenta), and αCHAT (red). White arrows point to examples of SYFP2+/αNos-C+ cells for both mouse and macaque panels. (**C**) Comparison of specificity of labeling in mouse and macaque based on IHC quantification. (**D**) Coronal view of the spatial distribution and abundance of SST-CHODL neurons in macaque Str. Navy blue dots represent SST-CHODL neurons shown over a grey background of all other cells for contrast. (**E-F**) Stereotaxic injection of AAV vector AiP15043 (best in class PVALB-PTHLH enhancer AiE1419m driving SYFP2) in mouse dorsal Str and macaque Cd. The total viral dose was 5E+8vg for mouse dorsal Str and 1.2E+11vg for macaque Cd. (**E**) Sagittal section of mouse brain showing DAPI, αGFP, and αPVALB staining. (**F**) Coronal section of macaque Cd region showing DAPI and αGFP staining. The image panels at right show magnified views of macaque Cd composite and individual channels with αGFP (green), PVALB (magenta) and PTHLH (red) RNA probes. White arrows point to examples of αGFP+/αPVALB+ cells in mouse and αGFP+/PVALB+/PTHLH+ cells in macaque. (**G**) Comparison of specificity of labeling in mouse and macaque based on IHC or mFISH quantification. (**H**) Coronal view of the spatial distribution and abundance of PTHLH-PVALB neurons in macaque Str. Red dots represent PTHLH-PVALB neurons shown over a grey background of all other cells for contrast. Abbreviations: Str, striatum; Cd, caudate.

SST-CHODL interneurons were robustly labeled by best-in-class enhancer AiE1426m in both mouse (**Fig. 3A**) and macaque (**Fig. 3B**). This enhancer drove reporter expression in neurons positive for the known SST-CHODL cell type marker, neuronal nitric oxide synthase (NOS), and negative for CHAT in both species (**Fig. 3B**), with SSv4 and IHC validation showing well matched on-target labeling of 81% in mouse and 77% in macaque, (**Fig. 3C**). Notably, the enhancer specificity by stereotaxic injection route in mouse dorsal striatum (81%) was lower than specificity measured by scRNA-seq following RO injection (99%), in agreement with prior routes of administration comparison data for this enhancer (Hunker et al. 2025), but it was essential to match delivery route for the most accurate comparison of mouse versus macaque enhancer-AAV specificity *in vivo*. In support of conserved high specificity, the morphological features of transduced cells—small somata, elongated wispy dendrites, and characteristic sparse arborizations (**Fig. 3B**)—recapitulated expected features for this class, as well as the overall distribution based on macaque BG spatial transcriptomics ground truth data (**Fig. 3D; ABC Atlas**).

For PTHLH-PVALB interneurons, the best-in-class enhancer AiE1419m produced specific labeling in both mouse (**Fig. 3E**) and macaque (**Fig. 3F**). Enhancer-AAV stereotaxic injection in mouse dorsal striatum produced highly specific labeling of PVALB-PTHLH interneurons (93% specificity by mFISH). In macaque, comparable mFISH analysis in the caudate injection site confirmed conserved specificity of enhancer-driven expression in PVALB-PTHLH interneurons with 90% specificity (**Fig. 3G**). Finally, the morphological features of transduced cells matched expected properties of PTHLH-PVALB interneurons–including medium-diameter soma and compact, multipolar dendrites (**Fig. 3E-F**)–while their medium-sparse distribution in dorsal striatum corresponded to that observed by spatial transcriptomics data (**Fig. 3H**).These results demonstrate that transcriptionally and epigenomically informed enhancer selection can successfully generalize to NHP, including for relatively sparse interneuron types.

### Enhancer AAVs for ventral striatum neuron types across species

The ventral striatum contains multiple highly distinctive Drd1(D1)-expressing cell types not captured by traditional dorsal MSN classifications. Two such populations—the D1-ICj granule cells of the and the D1-NUDAP cells of the ventral striatum—showed strong differential chromatin accessibility in our multi-species atlas and thus served as key targets for CERP prioritization. Here we provide a vignette comparing best-in-class enhancer performance for these distinctive ventral striatum neuron types in mouse versus macaque *in vivo*.

Differentially accessible peaks unique to each type yielded enhancer candidates with strong selective enrichment and clear separation between D1-ICj and D1-NUDAP chromatin states in both mouse and macaque (**Fig. 4A**). Consistent with spatial distribution patterns of these unique ventral striatum cell populations (**Fig. 4B**), mouse *in vivo* screening identified best-in-class enhancers with on-target specificity of 92% and 83% for D1-ICj granule cells (AiE1476q and AiE1223q, respectively; **Fig. 2C** and **Fig. 4C**) and 94% and 71% for D1-NUDAP (AiE1485q and AiE3010q, respectively; **Fig. 2C** and **Fig. 4D**) neurons by scRNA-seq analysis. We co-injected enhancer-AAVs for D1-ICj (AiE1476q) and D1-NUDAP (AiE3009q) and demonstrated duplex RO injection with mutually exclusive labeling in contrasting fluorophores (**Fig. 4E**). Off-target labeling outside the islands of Calleja or D1 NUDAP interface islands was minimal, consistent with the distinctive anatomical and neurochemical domains of these types.

**Figure 4:**
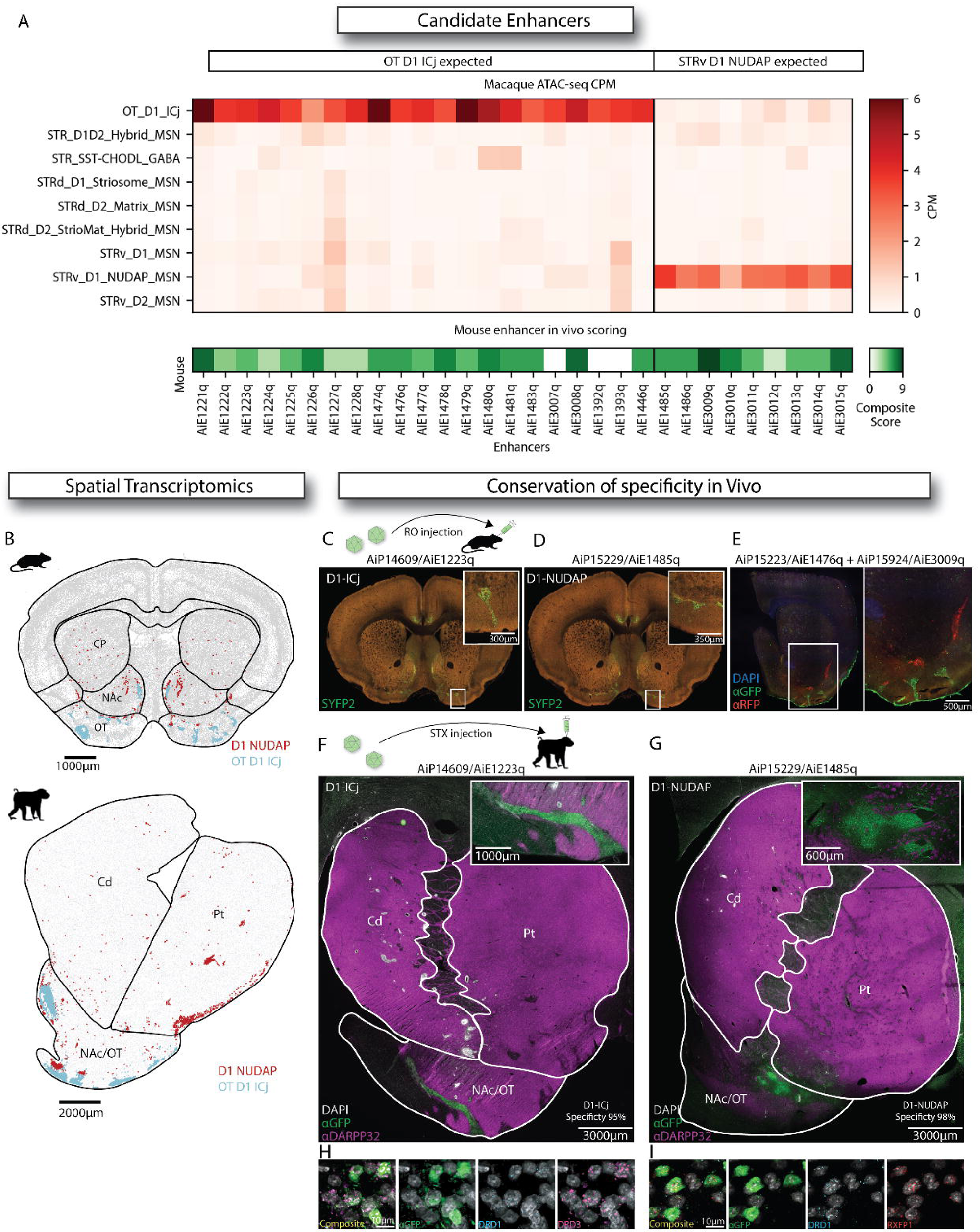
Cross-species comparison of ventral striatal cell type populations in mouse and macaque. (**A**) Heatmap of ATAC-seq counts per million (CPM) in-vivo scoring for candidate enhancers for OT D1-ICjs and STRv D1-NUDAP. (**B**) Coronal views of spatial transcriptomics data for OT D1-ICjs (blue) and STRv D1-NUDAP (red) for mouse (top) and macaque (bottom). (**C,D**) Coronal mouse brain sections showing native SYFP2 reporter expression following RO injections of D1-ICj and D1-NUDAP enhancer AAVs, respectively. Nissl stain (red), SYFP2 (green). (**E**) Coronal view of expressions in mouse RO duplex injection of D1-ICjs and STRv D1-NUDAP enhancers IHC. DAPI (blue), αGFP (green), αRFP (red). (**F,G**) Coronal view macaque stereotaxic injections of D1-ICjs and STRv D1-NUDAP enhancers IHC, respectively. DAPI (gray), αGFP (green), αDARPP32 (magenta). (**H,I**) Magnified tiles showing RNAscope in macaque for D1-ICjs and STRv D1-NUDAP enhancers using αGFP (green) Ab, DRD1 (cyan) and DRD3 (magenta) for D1-ICjs, and DRD1 and RXFP1 (red) probes for STRv D1-NUDAPs, respectively. All injections in macaque were performed using Brainsight (Rogue Research) surgical robot. Abbreviations: OT, olfactory tubercle; ICjs, Islands of Calleja; STRv, ventral striatum.

Cross-species evaluation in macaque revealed similarly striking enhancer activity, matching the expected spatial distribution for both D1-ICj (**Fig. 4F**) and D1-NUDAP (**Fig. 4G**) in the ventral striatum injection sites. RNAscope validation confirmed DRD1+/DRD3+ co-labeling in D1-ICj (**Fig. 4H**) and DRD1+/RXFP1+ co-labeling in D1 NUDAP (**Fig. 4I**) with on-target specificity of 95% and 98%, respectively, and consistent with prior delineation of maker genes for these neuron types in macaque ventral striatum (He et al. 2021). These data establish the first viral genetic tools to target D1-ICj and D1-NUDAP neurons in both rodents and primates. Their clustered spatial distribution and tight anatomical apposition in the ventral striatum but with unique transcriptomic identities make them ideal targets for future circuit dissection and behavioral studies.

### Pan-dopaminergic (DA) and DA subtype-selective enhancer AAVs

DA neurons of the substantia nigra pars compacta (SNc) and ventral tegmental area (VTA) comprise multiple transcriptionally distinct subtypes that differ in marker genes, projection targets, physiological profiles, and vulnerability in disease (Poulin et al. 2020). Analysis of spatial transcriptomics data in the mouse ABC atlas (Yao et al. 2023) revealed multiple spatially segregated DA neuron supertypes and clusters (subtypes) (**Fig. 5A-C**). These subtypes display differential gene expression along the dorsal–ventral axis, including known marker genes of DA neurons such as Tyrosine hydroxylase (*Th*), as well as by dorsal- (*Calb1*) and ventral-tier (*Aldh1a*) enriched marker genes (**Fig. 5D**). These molecular distinctions align with the defining axon projection targets in the striatum for each subtype, e.g., dorsal-tier DA neurons have axons projecting preferentially to ventromedial striatum, whereas, conversely, ventral-tier DA neurons have axons projecting predominantly to dorsolateral striatum (**Fig. 5E**).

**Figure 5.**
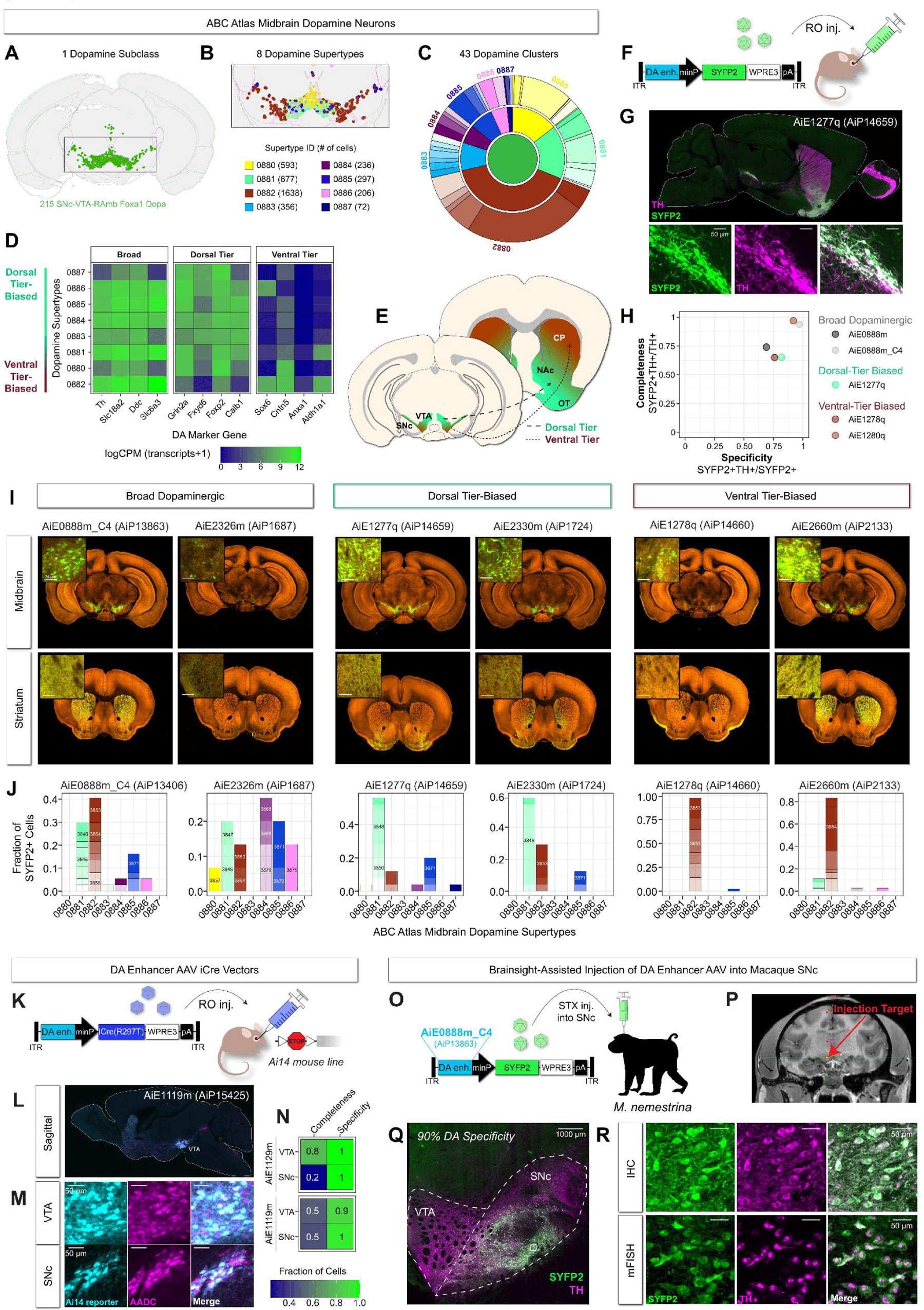
Enhancer-AAV targeting of midbrain dopaminergic neuron subtypes. (**A-C**) Spatial organization of midbrain dopaminergic populations from the ABC Atlas, shown at the levels of (**A**) DA neuron subclass, (**B)** DA neuron supertypes, and (**C**) DA neuron clusters. (**D**) Heatmap showing expression of selected dopaminergic marker genes across dopamine supertypes, organized to reflect broad, dorsal-, and ventral-tier associations. (**E**) Schematic illustrating canonical projection of dorsal- and ventral-tier dopaminergic neurons to their ventral and dorsal striatum targets, respectively. (**F**) Schematic of dopaminergic (DA) enhancer-AAV design and retro-orbital (RO) delivery in mouse. (**G**) Representative sagittal mouse brain section showing enhancer-driven SYFP2 expression in midbrain, with immunolabeling for tyrosine hydroxylase (TH). Bottom panels show higher-magnification views of SYFP2, TH, and merged channels. (**H**) Scatter plot depicting completeness versus specificity metrics for dorsal-tier, ventral-tier, and pan-DA enhancer-AAVs evaluated by single-cell RNA sequencing (SSv4). (**I**) Representative coronal sections of mouse midbrain and striatum showing SYFP2 expression driven by broad, dorsal-tier-biased, and ventral-tier-biased DA enhancers. Insets show higher-magnification views of labeled axons or somata. (**J**) Distribution of SYFP2-positive cells across ABC Atlas dopaminergic clusters for each enhancer, quantified by SSv4. (**K**) Schematic of DA enhancer-AAV-iCre vectors and RO delivery in Ai14 reporter mice. (**L**) Sagittal section showing Cre-dependent reporter expression following enhancer-AAV-iCre delivery. (**M**) High-magnification images of ventral tegmental area (VTA) and substantia nigra pars compacta (SNc) showing reporter expression and dopaminergic marker labeling. (**N**) Heatmap summarizing completeness and specificity metrics for Cre-dependent DA enhancer-AAVs in VTA and SNc. (**O**) Schematic of Brainsight-assisted stereotaxic injection of DA enhancer-AAV into macaque SNc. (**P**) Magnetic resonance image indicating injection target location in macaque midbrain. (**Q**) Representative midbrain section from macaque showing SYFP2 and TH immunolabeling following enhancer-AAV delivery. (**R**) High-magnification immunohistochemistry (IHC) and multiplexed fluorescence in situ hybridization (mFISH) images showing SYFP2 expression and dopaminergic marker labeling in macaque.

We screened 105 DA enhancer-AAV candidates with predicted activity in the SNc-VTA-RAmb Foxa1 Dopa subclass by RO injection in mice and focused subsequent detailed analysis below on 7 vectors with high performance. We identified enhancers that exhibited subtype selective labeling of dorsal-tier and ventral-tier midbrain DA neurons, as well as pan-dopaminergic enhancers with broad activity in the SNc-VTA-RAmb Foxa1 Dopa subclass. Two of each category were chosen for IHC and quantification with anti-TH antibody. For example, the best-in-class dorsal-tier DA subtype enhancer AiE1277q labeled neurons with axons projecting preferentially to ventromedial striatum and co-labeling with TH (**Fig. 5G**). Additional best-in-class DA enhancers showed high specificity and completeness of labeling in the IHC analysis using anti-TH staining (**Fig. 5H**). Whole brain imaging by serial two-photon tomography (STPT) for exemplary DA enhancers further enabled detailed evaluation of midbrain DA neuron soma labeling versus terminal axon projections in the striatum (**Fig 5I**). The matching set of exemplary pan-DA neuron and dorsal- and ventral-tier DA subtype enhancers were additionally evaluated by scRNA-seq with fine mapping at the transcriptomic supertype level in mouse (**Fig. 5J**). DA neuron enhancers AiE0888m_C4 and AiE2326m mapped to the broadest representation of DA neuron supertypes and are therefore classified as pan-DA targeting tools. Enhancers with preferential activity in dorsal-tier DA neurons (enhancers AiE1277q and AiE2330m) achieved 96% and 100% specificity at the SNc-VTA-RAmb Foxa1 Dopa subclass level, and showed strongest bias in mapping to finer supertypes 0881, 0882, and 0885. Conversely, enhancers labeling ventral-tier DA neurons—those disproportionately affected in Parkinson’s disease (Damier et al. 1999; Kamath et al. 2022)—achieved a remarkable 97.6% (AiE1278q) and 83.3% (AiE2660m) specificity of mapping to the high confidence ventral tier DA neuron mouse supertype 0882 (Fushiki et al. 2024), with little to no activity in other DA neuron supertypes. These results suggest a more nuanced relationship between anatomically-defined and transcriptomically-defined DA neuron types.

The DA neuron enhancers with highest midbrain DA neuron specificity and lowest off-target labeling across the brain were selected to make derivative iCre vectors and injected in Ai14 mice (**Fig. 5K**), enabling high-sensitivity Cre-dependent labeling for flexible intersectional strategies or other genetic perturbations. Enhancers AiE1129m and AiE1119m drove Cre-mediated recombination of tdTomato reporter expression with the expected spatial patterns and 90-100% DA neuron specificity in both SNc and VTA regions (**Fig. 5L-M**), albeit with variable completeness of labeling (**Fig. 5N**). Retroorbital virus doses were intentionally chosen to maximize specificity of midbrain DA neuron labeling while minimizing off-target recombination in the rest of the brain. Cross-species testing of broad DA neuron enhancer AiE0888m_C4 by stereotaxic injection targeted to SNc in macaque *in vivo* (**Fig. 5O-P**) revealed comparable levels of specificity for TH+ dopamine neurons (90%; **Fig. 5Q-R**) as observed in mice, with close agreement of specificity measurements by IHC and mFISH.

Together, these data demonstrate that enhancer-AAV strategies can be leveraged to selectively label the SNc-VTA-RAmb Foxa1 Dopa neuromodulatory neuron subclass spanning SNc and VTA regions. Remarkably, our screening also led to the discovery and validation of exemplary dorsal and ventral tier midbrain DA neuron subtype enhancers. This provides a critical foundation for mechanistic dissections of DA neuron subtype function, resilience, and degeneration in movement and affective disorders.

### Sequence and chromatin features predictive of BG enhancer performance *in vivo*

We performed a comprehensive analysis of open chromatin and sequence-derived features to characterize the 169 validated BG enhancers derived from the macaque genome. We examined a set of sequence and genomic features related to chromatin accessibility, evolutionary conservation, signal shape, gene specificity, distance to the nearest transcription start site, and GC content (**Table S2**). Within each track (**Fig. 6A**), higher feature values are displayed on the left and lower values on the right, allowing intuitive visual comparison of features. Together, these features capture both the chromatin and intrinsic sequence properties of candidate regulatory elements.

**Figure 6:**
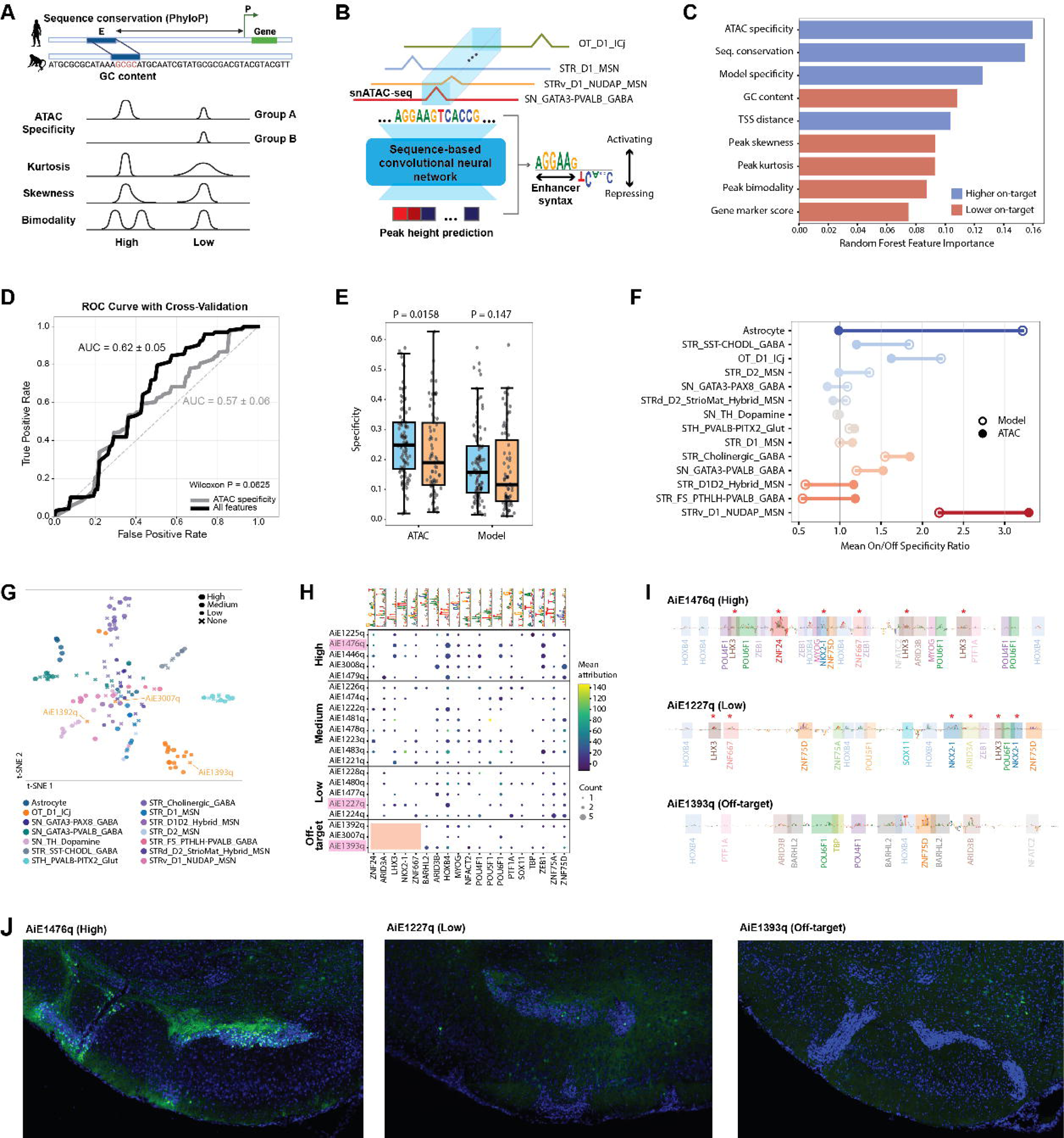
Sequence and chromatin features predictive of BG enhancer performance. (**A**) Overview of features collected for open chromatin data (E, enhancer; P, promoter). Higher feature values are shown on the left and lower values on the right. (**B**) CREsted model schematic. Pseudo-bulk group-aggregated BigWig tracks are used as input to predict peak height for each group and species. (**C**) Random forest feature-importance plot. Blue bars indicate features with higher mean values in on-target enhancers, and red bars indicate features with higher mean values in off-target enhancers. (**D**) AUC–ROC curves evaluating model performance using 5-fold cross-validation. The grey curve represents ATAC specificity alone, and the black curve represents all nine features from panel C. (**E**) Bar plots comparing on-target (blue) and off-target (red) enhancers for ATAC specificity (left) and CREsted specificity (right). (**F**) Scatter plot showing mean on/off ratios for ATAC and CREsted specificity per group. Filled circles denote ATAC specificity and open circles denote CREsted specificity. (**G**) t-SNE projection of CREsted predictions for all enhancers, colored by group. 3 OT_D1_ICj enhancers are labeled that had off-target activity. (**H**) Dot plot of seqlet cluster patterns matching BG atlas paper patterns annotated with JASPAR motifs using Pearson correlation, shown across OT_D1_ICj sequences ordered by specificity. Dot color indicates mean pattern attribution per sequence, and dot size indicates the number of patterns per sequence. Seqlets that were absent in off-target enhancers are highlighted with a rectangle. (**I**) Representative 3 of 21 OT_D1_ICj enhancers showing patterns present only in on-target enhancers (asterisks). (**J**) Epifluorescence images of on-target and off-target enhancers for the OT_D1_ICj group.

We leveraged DeepMacaqueBG (Johansen et al. 2025), a pre-trained CREsted (Kempynck et al. 2025) model of macaque BG cell types (**Fig. 6B**), to predict cell type-specific open chromatin based on DNA sequence of the 169 BG enhancers. We evaluated the relative importance of each chromatin and sequence feature in distinguishing on-target versus off-target enhancers using a random forest model (**Fig. 6C**). ATAC specificity was most predictive for functional enhancers (**Fig. S1**) as has been reported for neocortex (Johansen et al. 2025). On-target enhancers also had significantly higher sequence conservation (phyloP score) (**Fig. S2**) and higher DeepMacaqueBG sequence predictions of chromatin accessibility (**Fig. 6C**). Collectively, all features were more informative than ATAC-seq specificity alone at predicting enhancer activity (**Fig. 6D**). Next, we compared features of 154 enhancers for 14 groups with at least 1 on-target and 1 off-target enhancer. On average, ATAC and DeepMacaqueBG specificity was higher in on-target enhancers (**Fig. 6E**), and sequence differences captured by the model were more informative than ATAC specificity alone at identifying on-target enhancers for 6 out of 14 BG groups (**Fig. 6F**).

To investigate sequences contributing to enhancer function, we examined base-pair resolution predictions of enhancer activity. The DeepMacaqueBG embedding organized enhancer sequences into coherent clusters that defined Group specific sequence patterns (**Fig. 6G**). To move beyond prediction and toward mechanism by disentangling functional from non-functional enhancers, we examined the TF motifs used by the CREsted model to distinguish basal ganglia (BG) groups, focusing on group-specific patterns (Johansen et al. 2025) and testing whether these patterns are present in on-target enhancers using TF-MINDI. In the OT-D1-ICj group, we identified five DNA patterns, annotated with non-redundant vertebrate JASPAR motifs, that were present only in on-target enhancers and absent from off-target enhancers (**Fig. 6H–I**). These results indicate that functional enhancers are characterized by distinctive sets of motifs, and more patterns in enhancers with high and medium activity (**Figs. 6H–J** and **S3**).

Taken together, this retrospective analysis indicates that understanding functional enhancer activity in the BG requires a comprehensive multimodal approach that integrates chromatin-level features such as ATAC-seq specificity, evolutionary information such as phyloP conservation, and sequence-level grammar learned by deep models such as CREsted.

### Comparison of model prediction and *in vivo* performance of BG enhancer orthologs

Enhancers in the CERP-BG library were selected either from macaque or mouse chromatin accessibility data, raising a practical question for cross-species tool usage. We asked whether enhancer sequences identified in one species can be used interchangeably in another, or whether species-matched orthologs are required for optimal activity. Although transcription factor repertoires and trans-regulatory environments are largely conserved across mammals (Stergachis et al. 2014), we tested this assumption directly by comparing the *in vivo* activity of orthologous mouse, macaque, and human BG enhancer sequences in the mouse.

To this end, we cloned putative orthologs of best-in-class macaque enhancers from the human and mouse genomes and tested all three sequences—macaque (parent sequence), human ortholog, and mouse ortholog—under matched conditions in our standardized mouse RO screening pipeline (**Fig. 7A**). We performed this analysis for eight BG cell type targets spanning inhibitory, excitatory, and neuromodulatory populations, enabling an assessment of generality across diverse regulatory programs.

**Figure 7.**
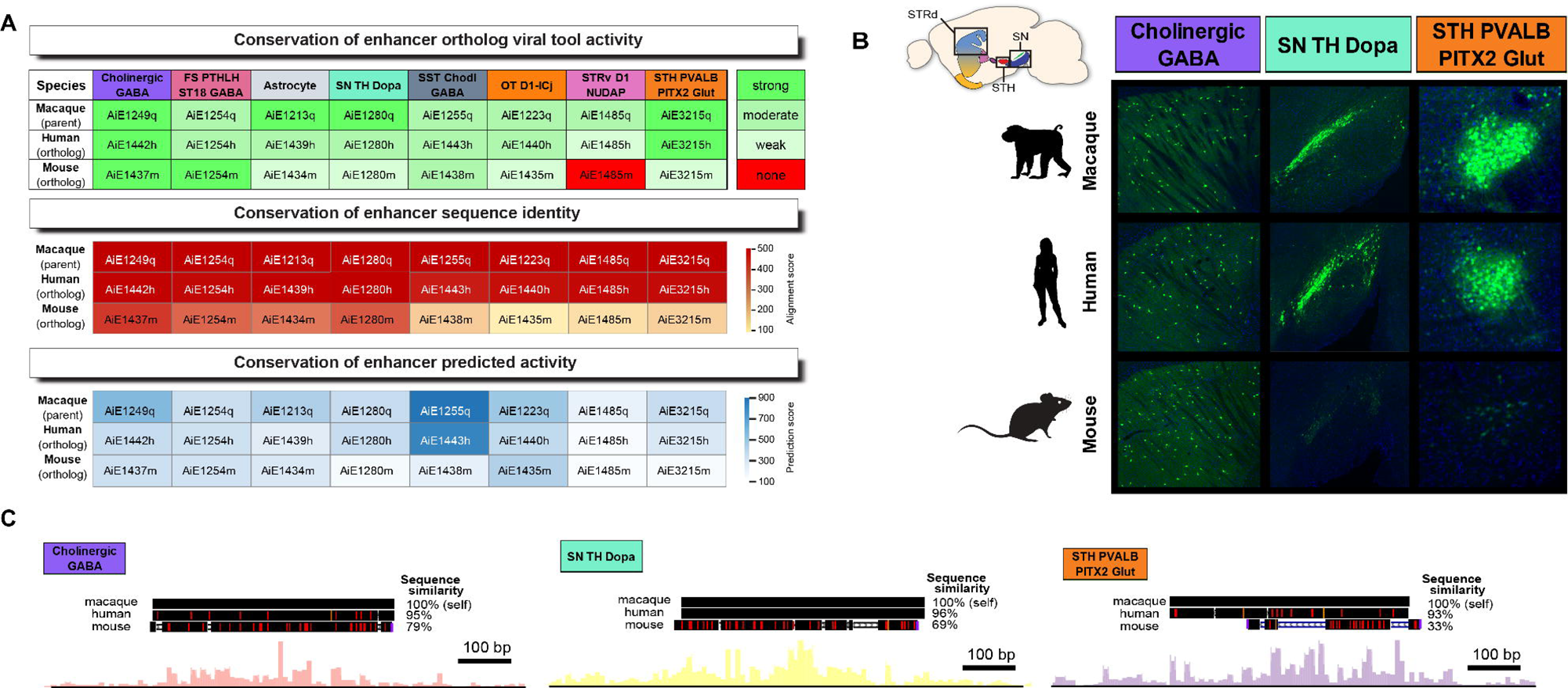
Cross-species conservation of enhancer sequence and viral tool activity. (**A**) Heatmaps comparing tested orthologs of best-in-class enhancers for 8 Groups. For each target, enhancer sequences identified in macaque were matched to putative human and mouse orthologs and evaluated using the standard mouse primary screening pipeline. Heatmaps summarize enhancer-AAV activity across species (top), sequence conservation based on BLAT alignment scores (middle), and predicted enhancer activity scores from a macaque-trained CREsted models (bottom). (**B**) Representative sagittal mouse brain sections showing SYFP2 expression driven by macaque, human, and mouse orthologs of selected enhancers targeting Str Cholinergic GABA, SN TH Dopa, and STH PVALB-PITX2 Glut neuron populations. Schematic insets indicate the targeted BG regions. (**C**) Genome browser views of representative enhancer loci showing orthologous sequence alignments across macaque, human, and mouse, with aligned conservation tracks and group-specific ATAC-seq chromatin accessibility.

Across most cell types, the specificity of enhancer activity was broadly conserved: human and mouse orthologs typically drove reporter expression in the same target populations as the original macaque-derived sequences. However, the strength and density of labeling in the target BG cell type varied substantially between orthologs, with the enhancer originally selected based on differential chromatin accessibility in macaque generally exhibiting the strongest and most specific activity in mouse BG. In one case—STRv D1 NUDAP—the mouse ortholog failed to drive detectable on-target labeling despite robust activity of the macaque and human sequences (**Fig. 7A,B**).

We asked whether these differences could be explained by primary sequence similarity. BLAT-based alignment scores showed the expected gradient of conservation, with human orthologs generally more similar to macaque (∼95% conservation) than mouse orthologs (∼70% conservation) (**Fig. 7A**, middle). However, sequence identity alone did not reliably predict functional outcomes: SST Chodl GABA and OT D1-ICj enhancers had relatively low macaque–mouse sequence conservation and retained similar activity, while Astrocyte and SN TH Dopa enhancers had relatively high sequence conservation and divergent activity.

We next evaluated whether sequence-based predictive models could account for ortholog performance. Applying the macaque-trained CREsted model to all orthologous sequences revealed partial concordance with experimental results. In particular, the STRv D1 NUDAP and STH_PVALB-PITX2_Glut mouse orthologs received low predicted activity scores, consistent with their failure or very weak activity *in vivo,* respectively (**Fig. 7A**, bottom). However, across the full set of enhancers, prediction scores alone did not fully capture the observed spectrum of ortholog performance, suggesting additional regulatory grammar.

Together, these results demonstrate that cross-species conservation of enhancer activity in the BG is not dictated by sequence identity alone, nor strictly by species-of-origin, but instead reflects preservation of essential cis-regulatory grammar. These findings underscore the importance of selecting enhancers based on cell type-resolved chromatin accessibility and regulatory motif composition, rather than defaulting to species-matched orthologs, when designing cross-species viral tools.

### Limitations of this study

Despite the breadth of this BG enhancer-AAV toolkit, additional enhancers will be required to access the full cellular diversity of the BG taxonomy. Our collection was designed to target broad classes and subclasses rather than the most granular cell types, as this represents the immediately actionable level of viral genetic access to serve the widest range of researchers in the neuroscience community–especially for research in NHP and other non-mouse models. It remains to be seen if differential chromatin accessibility or even AI-guided synthetic design can support efficient enhancer discovery for the finest transcriptomic brain cell types. Nonetheless, the validated BG enhancer collection presented here provides a strong foundation for iterative refinement. There are also notable challenges for applying these new tools for clinical applications where targeting of human BG cell types could be hugely impactful across a wide range of debilitating brain disorders involving BG dysfunction. Validation of enhancer activity in human BG remains largely infeasible, and current stem cell-derived and organoid or assembloid models do not yet recapitulate the full cellular diversity of the mature human brain or BG (Levy and Paşca 2023; Miura et al. 2024). The demonstrated conservation of enhancer activity from mouse to macaque shown here is very encouraging but does not guarantee similar performance in humans or in specific disease contexts. Thus, while this toolkit represents a substantial advance for research in cellular and systems neuroscience, there remain important hurdles to clear on the path to future cell type-targeted AAV gene therapies.

## DISCUSSION

The BG sit at the center of motor control, reinforcement learning, and action selection, and their dysfunction underlies a spectrum of neurological disorders. Yet, for all their clinical and conceptual importance, the BG have remained disproportionately difficult to interrogate at cellular resolution, particularly beyond the mouse model. Given the central importance of NHPs including marmoset and macaque in neuroscience research globally (Okano 2021; Tremblay et al. 2020) and their value for understanding cellular diversity and function that are closer to humans than in rodents (Bakken et al. 2021; Chen et al. 2025), it is essential to fill this gap and level-set capabilities for cell type-specific genetic access and perturbation across species. In this study we lay out a new framework for brain cell type enhancer discovery and cross-species validation that explicitly incorporates a cross-species consensus BG taxonomy (Johansen et al. 2025) with a cross-species enhancer ranking pipeline and *in vivo* functional validation in both rodent and primate to deliver a comprehensive enhancer-AAV toolkit spanning the principal input, relay, output, and neuromodulatory nodes of BG circuitry. By anchoring our end-to-end enhancer discovery and validation process in this consensus cross-species taxonomy we avoid confusion in aligning cellular taxonomies arising from disparate sources in different species and with differing nomenclature. By design, these tools hold great promise for enabling genetic access to homologous cell types in various other mammalian model organisms as well as for possible human clinical applications.

The present study directly expands our prior work on enhancer-AAV vectors for targeting canonical cell types of the dorsal striatum including D1 and D2 MSNs, and local interneuron subclasses SST-CHODL, PVALB-PTHLH, and Cholinergic neurons (Hunker et al. 2025). We folded in ventral striatum specificity data to explicitly reveal whole striatum specificity of the best-in-class striatal enhancers, revealing enhancers with highest specificity in each region, respectively. Next, we identified and validated enhancer-AAV vectors for targeting additional ventral striatum cell types, with emphasis on historically less well-studied and non-canonical types including D1/D2 hybrid MSNs, STRv D1 NUDAP and D1-ICj granule cell type. Established BG circuit frameworks anchored around canonical direct (D1 MSN) and indirect (D2 MSN) striatal projection pathways (Albin et al. 1989; DeLong 1990) do not capture the full complement of these specialized ventral striatum cell types that occupy anatomically restricted niches (He et al. 2021; Andraka et al. 2024) and have emerging critical roles in motor control, habitual behavior, and affect (Castro and Bruchas 2019; Zhang et al. 2021; Zhang et al. 2023). In particular, D1-ICj granule cells have been the subject of recent functional interrogation studies in mice, and offer a compelling entry point into ventral striatal microcircuits and clear example of a cell type with emerging links to stereotyped behaviors such as grooming and potential relevance to compulsive phenotypes (Zhang et al. 2021). Genetic targeting and perturbation of D1/D2 hybrid MSNs and STRv D1 NUDAP cells has been far more challenging to date. Interestingly, D1/D2 hybrid MSNs and STRv D1 NUDAP cells have similar transcriptomic signatures and partially overlapping spatial distributions and are therefore often grouped in the literature under broader descriptors such as eccentric MSNs (also known as eSPNs) (Johansen et al. 2025; Gayden et al. 2023; Saunders et al. 2018). These types illustrate a recurring theme: as atlases sharpen cell type distinctions, the experimental bottleneck shifts from establishing cell type existence to developing tools for genetic perturbation and cross-species interrogation. The tools reported here begin to make that shift tractable. By contrast, D1-NUDAP neurons remain comparatively enigmatic. From a practical perspective, this asymmetry is precisely where tools have maximal leverage: genetically precise access turns an atlas-defined population into a foothold for testable hypotheses aimed at unraveling the role of a cell type in circuit function.

Among the BG populations highlighted here, midbrain DA neuron subtypes illustrate the clinical and conceptual value of resolving fine-grained diversity. Although there are existing viral genetic tools with cell type promoters and enhancers to broadly target midbrain DA neurons in rodents and NHPs (Caplan et al. 2025; Oh et al. 2009), no such enhancers exist for selective labeling of midbrain DA neuron subtypes. Instead, researchers have relied on retrograde (Tervo et al. 2016) or intersectional (e.g., Cre-dependent) viral labeling methods (Stauffer et al. 2016) that suffer from low efficiency or high complexity. Importantly, dorsal- and ventral-tier DA neurons differ in spatial organization, axon projection patterns, molecular profiles, and vulnerability to degeneration in Parkinson’s disease (Pereira Luppi et al. 2021; Kamath et al. 2022). The tools developed here for viral genetic targeting of dorsal- and ventral tier DA neuron subtypes simplify the experimental paradigm and will accelerate progress in deciphering the relative contribution of DA neuron subtypes to specific behaviors or computations, and which subtype properties predict degeneration or resilience in disease. It may now be possible to achieve biased delivery of therapeutic payloads to the most vulnerable ventral tier DA neurons while avoiding resilient types. A clear opportunity for future work is to increase subtype resolution further, moving from tier-biased groups to finer transcriptomic and spatial types. Doing so will likely require larger training sets of validated enhancers per subtype, improved sequence models that better capture combinatorial grammar, and experimental paradigms that integrate labeling with projection mapping and perturbation. The same roadmap–starting with broad class and subclass enhancers and progressing to fine subtype enhancer discovery–can naturally be applied to develop novel viral genetic tools to access other neuromodulatory systems in the brain.

The enhancer-AAVs for targeting STN neurons provide a complementary translational bridge. The STN is a prime target for deep brain stimulation (DBS) in Parkinson’s disease (Okun 2012) and may hold promise in treating obsessive-compulsive disorder (Mallet et al. 2008), and its position in BG circuitry makes it an appealing node for both basic mechanistic dissection and therapeutic modulation. Interestingly, similar to optogenetic perturbation of D1-ICj granule cells (Zhang et al. 2021), activation of STN neuron populations in mice elicited grooming behavior, whereas chronic silencing suppressed it, likely through altering plasticity in BG circuits (Parolari et al. 2021). The new STN neuron specific enhancer-AAVs should facilitate similar mechanistic studies into DBS and the circuitry basis of motor-related behaviors beyond the mouse model. Remarkably, some exemplary STN enhancers showed striking whole-brain specificity following systemic virus administration in mice, suggesting that enhancer-AAV delivery can achieve the kind of anatomical restriction of cargo expression that is often assumed to require invasive (e.g., stereotaxic injection) targeting. This raises an exciting, if still speculative, possibility: enhancers with exceptional whole brain specificity could enable more precise and potentially less invasive therapeutic interventions than current approaches. The empirical demonstration of enhancers with dominant activity in just one cell type in the whole brain has important implications for future efforts in synthetic enhancer design, understanding cell type enhancer regulatory logic, and suggests that further integration of brain and body wide single cell epigenetic datasets will hold exceptional value for improving enhancer discovery into the future.

Beyond adding novel tools for BG cell type genetic access, this work also clarifies the functional mechanisms that determine enhancer cell type specificity and enhancer-AAV success. Our modeling supports a multimodal view in which cell type-resolved chromatin accessibility is a dominant predictor, but not sufficient on its own. Adding sequence-level predictions and evolutionary constraints improves separation of on-target and off-target candidates, indicating that ultimate enhancer performance in vivo reflects both local epigenomic context and intrinsic cis-regulatory grammar (Kempynck et al. 2025). Seqlet- and motif-level analyses further suggest that some BG groups rely on distinctive TF-site combinations enriched in on-target enhancers. Although the number of experimentally validated enhancers remains limited, it was reassuring that group-specific sequence patterns identified as statistically enriched were preferentially present in functional enhancers, and more strongly represented in on-target compared to off-target examples. This concordance supports the biological relevance of the learned patterns, even though the signals are not uniform across all groups. The study therefore complements brain cell type atlases (Johansen et al. 2025) with a curated, functionally tested subset linking sequence features to in vivo performance. As the collection of validated enhancers grows, future work will expand these datasets and incorporate systematic in silico mutagenesis experiments to directly test the functional contribution of the identified motifs, helping to distinguish which transcription factor combinations are necessary versus sufficient for cell type specific activity.

Our enhancer ortholog analysis covering three species and eight different BG cell types underscores a related point about regulatory grammar versus raw sequence identity. In practice, cross-species tool use is often framed as a choice between species-matched orthologs and non-native mammalian enhancers, but the data argue against simple heuristics. Many enhancers retain specificity across human, macaque, and mouse despite substantial DNA sequence divergence, consistent with preserved key motifs, whereas some orthologs fail despite high overall similarity, consistent with disruption of critical regulatory features. Across all tests, primate enhancers identified from primate multiomic data performed comparably or better than mouse orthologs in mice, and the reverse was never observed. Accordingly, enhancer selection should proceed based on differential accessibility analysis using the highest quality of cell type-resolved open-chromatin data, with motif composition and learned sequence features as additional filters, rather than by species matching alone. These results also motivate broader ATAC-seq profiling across mammalian species to expand the pool of cell type-specific enhancer candidates.

Taken together, this study establishes a comprehensive, cross-species enhancer-AAV toolkit for the BG and a scalable workflow for extending such resources to other regions and species. By unifying atlas-grounded cell type definitions with high-throughput functional screening, ortholog comparisons, and sequence-informed modeling, we aim to shift enhancer-driven targeting from artisanal reagent development to a more systematic, interpretable discipline. The immediate payoff is practical—high-specificity viral tools to access BG cell types that have been difficult to target and manipulate, especially in NHPs. The longer-term payoff is mechanistic and predictive: validated enhancers connect cell type taxonomies to circuit hypotheses and reveal the cis-regulatory logic underlying neuronal identity across mammals.

## RESOURCE AVAILABILITY

### Lead contact

Requests for further information and resources should be directed to and will be fulfilled by the lead contact, Dr. Jonathan T. Ting.

### Materials availability

Plasmids generated in this study will be deposited to Addgene for distribution (in progress). Full plasmid DNA sequences and maps are available for each vector. Plasmids can be ordered through Addgene and require a standard UBMTA. There are no restrictions to plasmid availability for academic or not-for-profit research use. Commercial use requires licensing through the Allen Institute Office of Innovation.

### Data and code availability

⍰ scRNA-seq data have been deposited at Neuroscience Multi-omic Archive (NeMO): nemo:dat-ni6b3nk and will be publicly available at the date of publication at https://assets.nemoarchive.org/dat-ni6b3nk.
⍰ Primary screen EPI and STPT data have been deposited at Brain Image Library (BIL) and will be publicly available at the date of publication at https://doi.org/10.35077/g.1193.
⍰ Primary screen EPI and STPT data and metadata for many Tools described in this paper are available through an Allen Institute public web resource, the Genetic Tools Atlas (RRID:SCR_025643; https://portal.brain-map.org/genetic-tools/genetic-tools-atlas)
⍰ PeakRankR has been deposited at https://github.com/AllenInstitute/PeakRankR.
⍰ All original code has been deposited at https://github.com/AllenInstitute/PyPeakRankR and https://github.com/AllenInstitute/BG_CERP.
⍰ Any additional information required to reanalyze the data reported in this paper is available from the lead contact upon request.

## ACKNOWLEDGMENTS

This publication was supported by and coordinated through the BRAIN Initiative Armamentarium Consortium for Precision Brain Cell Access grant UF1MH128339 (B.P.L., B.Ta., G.D.H., J.T.T., T.E.B., T.L.D., and Y.K.) and the BRAIN Initiative Cell Atlas Network (BICAN) Consortium grant UM1MH130981 (E.S.L. and H.Z.). This work was also supported by the Allen Institute for Brain Science and the Research Foundation Flanders (FWO) PhD fellowship 1SH6J24N & V428025N (N.K.). The authors thank the founder of the Allen Institute, Paul G. Allen, for his vision, encouragement and support. We also thank the many support teams of the Allen Institute for Brain Science, as well as veterinary, husbandry, and surgery support staff of the WaNPRC. We thank Christopher English, Britni Curtis, and Jesse Day for support with various NHP experiment logistics. The WaNPRC is supported by the NIH Office of Research Infrastructure Programs (ORIP) under awards P51OD010425 and U420D011123. The contents of this study are solely the responsibility of the authors and do not necessarily represent the official view of the NIH, WaNPRC, or ORIP. We thank the University of Washington DISC lab for macaque MRI support.

## AUTHOR CONTRIBUTIONS

Conceptualization: BPL, BTa, JTT, TEB, TLD

Methodology: ACH, BPL, BTa, ET, GDH, JA, JTT, MEW, MHo, MNL, NJJ, NL, NT, NW, SSo, TEB, TLD, VO, WDL, YG, YK

Software: AO, MEW, MHo, NJJ, NK, SSo

Validation: ACH, GDH, JTT, MEW, MNL, NJJ, NT, SSo, TEB, WDL

Formal Analysis: ACH, BPL, DD, ET, JKM, JTT, MEW, MHo, MNL, MTS, NJJ, NK, NL, NT, SSo, TEB, VO, WDL, YB, YG

Investigation: AAO, AAy, ACH, AHua, APA, AR, BPL, CAP, CR, DB, DH, DJ, EL, ET, GDH, JA, JTT, JWa, LP, MB, MEW, MHo, MJT, MNL, MT, NID, NJJ, NN, NP, NT, NVS, NW, SDH, SL, SR, SSo, TCE, TCar, TEB, TEO, VO, WDL, YB, YG, YK, ZCJ

Resources: AAm, AO, AW, BO, BPL, BTa, BTh, CG, DN, DN, EG, GDH, HL, IE, JK, JR, JTT, JWi, KN, LS, MEW, MHo, MP, NJJ, RAM, RK, RN, SK, SSo, SW, SY, TEB, TJ, TLD, WHa, XO, YK

Data Curation: ACH, AO, BPL, BTa, BTh, DD, DR, EG, ET, GDH, JGo, JR, JTT, KAS, MEW, MHo, MNL, MT, NJJ, NT, NW, RAM, SSo, SW, TEB, TLD, VO, WDL, XO, YB

Writing - Original Draft: MEW

Writing - Reviewing & Editing: ACH, JTT, MEW, MNL, SSo, TEB, WDL

Visualization: ACH, JTT, MEW, MHe, MNL, SSe, SSo, TEB, WDL

Supervision: ABC, BPL, BTa, ESL, GDH, HZ, JA, JKM, JMil, JTT, KAS, LP, MJT, ND, NT, NVS, RAM, SDH, SSe, TEB, TLD, YK

Project Administration: ACH, BPL, BTa, BTh, GDH, JTT, MEW, MJT, SW, TEB, TLD, YK

Funding Acquisition: BPL, BTa, JTT, TEB, TLD

## DECLARATION OF GENERATIVE AI AND AI-ASSISTED TECHNOLOGIES

During the preparation of this work, the authors used OpenAI ChatGPT-5 to support certain coding tasks, assist with initial drafts of the main text to improve conciseness and readability, and identify potentially relevant citations. After using this tool, the authors reviewed and edited the content as needed and take full responsibility for the final content of the publication.

## SUPPLEMENTAL INFORMATION

**Figure S1, related to Figure 6. Column-normalized model specificity heatmap across BG cell types.** Rows represent cell types and columns represent ATAC peaks, ordered by group and functional status. Top annotation bars indicate group and status (high, medium, low, none). Colors show column-normalized values (0–1), with yellow indicating the strongest CREsted prediction signal per peak.

**Figure S2, related to Figure 6. Comparison of sequence and regulatory features between on-target and off-target macaque enhancers.** Boxplots show distributions of ATAC-seq specificity, PhyloP conservation, CREsted specificity, GC content, TSS distance, skewness, kurtosis, bimodality, and gene τ score. Each point represents an individual enhancer. P-values (Wilcoxon rank-sum test) are shown above each panel; * indicates statistical significance (Wilcoxon test, ATAC-seq specificity p = 0.01583, PhyloP = 0.001141).

**Figure S3, related to Figure 6. Pattern diversity across macaque enhancer activity classes.** Boxplots show the distribution of the number of total patterns per enhancer for high, medium, and low activity enhancers. Each point represents an individual enhancer. P-values from pairwise Wilcoxon rank-sum tests are shown above brackets. Enhancers with zero patterns did not contain any seqlets with an attribution score greater than 3.

**Table S1, related to Figure 1. Summary of 531 tested enhancer sequences for BG cell types and best-in-class tools presented in the paper.**

**Table S2, related to Figure 6. List of sequence and regulatory features values for 169 macaque enhancers.**

## METHODS

### Experimental model details Animals

All procedures involving mice were approved by the IACUC at the Allen Institute for Brain Science (protocols 2004, 2105, 2301, 2306, 2406, and 2010). Mouse tissue was obtained from 4 to 12-week-old male and female pure C57Bl6/J mice and adult Ai14 heterozygous mice. Animals were provided food and water *ad libitum* and were maintained on a regular 12-h day/night cycle with no more than five adult animals per cage.

### Enhancer selection and genomics analyses

To systematically identify cell type–specific enhancers across basal ganglia (BG) nuclei and species, we developed the Cross-species Enhancer Ranking Pipeline (CERP), an end-to-end computational and experimental framework that links atlas-defined cell types to prioritized candidate enhancers for *in vivo* validation (Figure 1B). CERP begins with cell type definition based on single-nucleus RNA sequencing (snRNA-seq) atlases generated from macaque (Johansen et al. 2025) or mouse (Yao et al. 2023) BG nuclei. Cell types were annotated according to atlas-resolved clustering and harmonized across species using the HMBA consensus nomenclature (Johansen et al. 2025), enabling cross-species alignment of transcriptionally homologous populations. For each annotated cell type, corresponding single-nucleus ATAC-seq (snATAC-seq) data were used to identify open chromatin regions enriched in that population.

The pool of candidate regulatory elements prioritized by CERP was populated using multiple complementary approaches. The primary source consisted of cell type-specific peaks identified through genome-wide differential accessibility analysis, retaining peaks meeting the following criteria: (i) significant enrichment in the target cell type relative to all other BG cell types (fold-change enrichment > 1.5, FDR < 0.05), assessed using MACS2; (ii) minimum accessibility strength exceeding 50 overlapping reads; and (iii) minimal off-target ATAC signal in other cell types, as assessed by genome browser inspection. These thresholds were relaxed for rare populations where even the most specific natural enhancers did not meet all criteria. In parallel, we identified additional candidates through targeted searches near established marker gene loci for major BG neuron classes using both our in-house multiome data and publicly available snATAC-seq resources, including the Cis-element Atlas (CATlas (Li et al. 2021)). Differentially accessible peaks within approximately 500 kb of BG cell type marker genes were manually inspected in genome browsers loaded with cell type-resolved accessibility tracks, enabling identification of candidates with strong target-population signal that may have been missed by automated differential peak calling. Previously characterized dorsal striatum enhancers included for ventral striatum characterization in this study were discovered as originally indicated in (Hunker et al. 2025). Finally, enhancers screened for other brain regions that exhibited cell type specific activity in BG were included where relevant. All candidate peaks, regardless of source, were subsequently filtered to exclude promoter-proximal regions (defined as within 2 kb of an NCBI RefSeq-annotated transcription start site) and to require evidence of cross-species sequence conservation via BLAT alignment between mouse and macaque, with preference given to peaks showing conserved chromatin accessibility in both species.

To move beyond simple fold-change ranking and integrate multiple predictive features, we developed PypeakRankr, a Python package for systematic collection and weighting of peak-level metrics from group-aggregated bigWig files. For each candidate enhancer, PypeakRankr computed chromatin accessibility specificity, evolutionary conservation (PhyloP), distance to the nearest TSS, signal distribution shape (kurtosis, skewness, bimodality), and sequence composition (GC content). These features were then combined to generate composite rankings that outperformed conventional approaches based on differential accessibility fold-change alone, allowing prioritization of candidates most likely to drive specific expression *in vivo*.

### Enhancer cloning into AAV vectors

Enhancers were cloned from human or C57Bl/6J mouse genomic DNA using enhancer-specific primers and Phusion high-fidelity polymerase (M0530S; NEB). Individual enhancers were then inserted into a recombinant single-stranded pAAV backbone that contained the beta-globin minimal promoter, fluorescent reporter gene (typically SYFP2), minimal woodchuck hepatitis posttranscriptional regulatory element (WPRE3), and bovine growth hormone polyA using standard molecular cloning as previously described (Graybuck et al. 2021; Mich et al. 2021). Plasmid integrity was verified via Sanger DNA sequencing and in some cases restriction digestion with agarose gel electrophoresis to confirm intact inverted terminal repeats (ITRs). In some cases, synthetic enhancer sequences such as concatenated cores were gene synthesized and subcloned into AAV vectors using standard restriction enzyme digestion and ligation. All AAV plasmids were propagated in NEB stable E.coli at 30°C growth condition to prevent spurious DNA rearrangements.

### AAV packaging and titer determination

Small-scale crude AAV preps were generated by triple transfecting 15 μg ITR plasmid, 15 μg AAV capsid plasmid, and 30 μg pHelper (Cell Biolabs) into one 15-cm plate of confluent HEK-293T cells using PEI Max (Polysciences Inc., catalog # 24765-1). At one day post-transfection we changed the medium to low serum (1% FBS), and after 3 days the cells and supernatant were collected, freeze-thawed 3x to release AAV particles, treated with benzonase nuclease (MilliporeSigma catalog #E8263-25KU) for 1 h to degrade free DNA, then clarified (3000 g × 10 min) and concentrated to approximately 150 μL by using an Amicon Ultra-15 centrifugal filter unit (NMWL 100 kDa, Sigma #Z740210-24EA) at 5000g for 30–60 min, yielding a titer of approximately 1.0E+13 to 1.0E+14 vg/mL. An efficient and cost-effective protocol for small scale crude AAV preparation has been shared on protocols.io (https://www.protocols.io/view/production-of-crude-aav-virus-extract-yxmvmx7r9l3p/v6) by the Allen Institute team. This protocol typically yields 5E+12vg from one 15 cm plate of AAV293 cells using the conventional adherent cell culture and triple transfection method with serotype PHP.eB.

For large-scale gradient preps, we transfected 10 x 15-cm plates of cells and purified by iodixanol gradient centrifugation. For measuring virus titers, we used ddPCR (Bio Rad; QX 200 Droplet Digital PCR System). We used primers against AAV2 ITR for amplification. Seven serial dilutions with the factor of 10 ranging from 2.5×10−2 to 2.5×10−8 were used for the measurement. Serial dilutions of 2.5×10−5 to 2.5×10−8 were used for fitting the dynamic linear range. Viral titer was calculated by averaging virus concentration of two dilutions within the dynamic linear range. A positive control of a known viral titer, and a negative control with no virus was also run along with all the samples.

### Retro-orbital (RO) and stereotaxic (STX) injection delivery of AAV vectors

Adult C57Bl6/J mice were briefly anesthetized by isoflurane anesthesia and injected with crude PHP.eB39 serotyped AAV virus at a dose range of 1E+10 to 1E+12vg into the retroorbital sinus of one eye in a maximum volume of 100μL. Virus stocks were diluted in sterile 1X phosphate buffered saline (1XPBS) solution as needed to achieve the intended dose and volume. For initial screening with enhancer-AAVs driving SYFP2, we routinely used 5E+11vg as the standardized RO dose. For stereotaxic injection surgery, C57Bl/6J, adult mice were deeply anesthetized to a surgical plane using an isoflurane vaporizer and placed into the stereotaxic injection frame. AAV virus was injected bilaterally into the dorsal striatum region (dSTR) using the following coordinates (in mm) relative to bregma: anterior/posterior (A/P) 0.8, medial/lateral (M/L) ±1.8 to 2.0, and dorsal/ventral (D/V) 2.6 to 3.0. A total volume of 500 nL containing 1E+12 to 1E+13 vg/mL virus stock was delivered at a rate of 50 nL per pulse with a Nanoject II pressure injection system. Before incision, the animal was injected with Bupivacaine (2–6 mg/kg) and post injection, the animal was injected with ketofen (2–5 mg/kg) and Lactated Ringer’s Solution; LRS (up to 1 mL) to provide analgesia. Mice that underwent STX injections were euthanized after 3–5 weeks post injection, transcardially perfused with 1XPBS followed by 4% paraformaldehyde (PFA), and the brains were dissected for further analysis.

### Tissue processing for slide-based epifluorescence imaging

Mice were anesthetized with isoflurane and perfused transcardially with 10 mL of 0.9% saline, followed by 50 mL of 4% PFA. The brain was removed, bisected along the midsagittal plane, placed in 4% PFA overnight and subsequently moved to a 30% sucrose solution until sectioning. From the left hemisphere, 30 μm sections were obtained along the entire mediolateral axis using a freezing, sliding microtome. Five sagittal ROIs, roughly 0.5, 1, 1.5, 2.3 and 3.5 mm from the midline, were collected and stained with DAPI (4′,6-diamidino-2-phenylindole dihydrochloride) and/or propidium iodide (PI) to label nuclei and to reveal cellular profiles, respectively. Stained tissue sections were slide mounted using Vectashield hardset mounting medium (Vector Laboratories, catalog # H-1400-10) and allowed to dry for 24 h protected from light. Once the mounting medium hardened, the slides were scanned with Aperio VERSA (Leica) or VS200 (Olympus) epifluorescence microscopes in the UV, green, and red channels, illuminated with a metal halide lamp. After passing QC, digitized images were analyzed by manual scoring of virus-mediated fluorescent protein expression throughout the brain with emphasis on striatal regions including caudoputamen, nucleus accumbens, and olfactory tubercle. Images were not acquired under matched conditions and have been adjusted to optimally highlight brain wide expression patterns for enhancer specificity comparison purposes.

### Serial two-photon tomography (Tissuecyte)

Mice were perfused with 4% PFA. Brains were dissected and post-fixed in 4% PFA at room temperature for 3–6 h and then overnight at 4°C. Brains were then rinsed briefly with PBS and stored in PBS with 0.01% sodium azide before proceeding to the next step. Agarose was used to embed the brain in a semisolid matrix for serial imaging. After removing residual moisture on the surface with a Kimwipe, the brain was placed in a 4.5% oxidized agarose solution made by stirring 10 mM NaIO4 in agarose, transferred through phosphate buffer and embedded in a grid-lined embedding mold to standardize its placement in an aligned coordinate space. The agarose block was then left at room temperature for 20 min to allow solidification. Brain tissue was additionally supported by generating a polyacrylamide network throughout the agarose block and spanning the brain-agarose interface. The agarose block was first left at 4°C overnight in a solution of 4.5% Surecast (Acrylamide:Bis-acrylamide ratio of 29:1) with 0.5% VA-044 activator, diluted in PBS. Agarose blocks were then placed back into new embedding molds containing a small amount of acrylamide solution and the top surface covered with parafilm (to reduce exposure to oxygen). Finally, specimens are baked for 2 h at 40°C and then stored in PBS with 0.1% sodium azide at 4°C until ready to image. The agarose block was then mounted on a 1 × 3 glass slide using Loctite 404 glue and prepared immediately for serial imaging.

Image acquisition was accomplished through serial two-photon (STP) tomography using six TissueCyte 1000 systems (TissueVision, Cambridge, MA) coupled with Mai Tai HP DeepSee lasers (Spectra Physics, Santa Clara, CA). The mounted specimen was fixed through a magnet to the metal plate in the center of the cutting bath filled with degassed, room temperature PBS with 0.1% sodium azide. A new blade was used for each brain on the vibratome and aligned to be parallel to the dorsoventral axis. Brains were imaged from the caudal end. We optimized the imaging conditions for both high-throughput data acquisition and detection of single axon fibers throughout the brain with high resolution and maximal sensitivity. The specimen was illuminated with 925 nm (EGFP, tdTomato, dTomato data) or 970 nm (SYFP2 data) wavelength light through a Zeiss ×20 water immersion objective (NA = 1.0), with 250 mW light power at objective. The two-photon images for red, green and blue channels were taken at 75 μm below the cutting surface. This depth was found optimal as it is deep enough to avoid any major groove on the cutting surface caused by vibratome sectioning but shallow enough to retain sufficient photons for high contrast images. In order to scan a full tissue section, individual tile images were acquired, and the entire stage was moved between each tile. After an entire section was imaged, the x and y stages moved the specimen to the vibratome, which cut a 100-μm section, and returned the specimen to the objective for imaging of the next plane. The blade vibrated at 60 Hz and the stage moved toward the blade at 0.5 mm per sec during cutting. Images from 140 sections were collected to cover the full range of mouse brain. It takes about 18.5 h to image a brain at an x,y resolution of ∼0.35 μm per pixel, amounting to ∼750 GB worth of images per brain. Upon completion of imaging, sections were retrieved from the cutting bath and stored in PBS with 0.1% sodium azide at 4°C.

### Immunohistochemistry

Brain slices were fixed in 4% PFA in phosphate buffered saline (PBS) at 4°C overnight or up to 48 h and then transferred to 1XPBS with 0.01% sodium azide as a preservative. Fixed slices were thoroughly washed with PBS to remove residual fixative, then blocked for 1 h at room temperature in 1XPBS containing 5% normal goat serum and 0.2% Triton X-100. After blocking, slices were incubated overnight at 4°C in blocking buffer containing one or more of the following primary antibodies: chicken anti-GFP (Aves, 1:1000 or 1:2000), rabbit anti-red fluorescent protein (Rockland, 1:500), mouse anti-ChAT IgG2b (Atlas Labs, 1:1000), Rabbit anti-nNos (Immunostar, 1:1000), Rabbit anti-Pvalb (Swant, 1:2500). Following the overnight incubation, slices were washed for 15 min three times with 1XPBS and then incubated for 2 h at room temperature in dye-conjugated secondary antibodies (1:1000; Invitrogen, Grand Island, NY) including BV480 (goat anti-rat)Alexa Fluor 488 (goat anti-chicken), Alexa Fluor 555 (goat anti-rabbit and goat anti-mouse), and Alexa Fluor 647 (goat anti-mouse). Slices were washed for 15 min three times with 1XPBS, followed by 5 μg/mL DAPI nuclear staining for 15 min. The slices were then dried on glass microscope slides and mounted with Fluoromount-G.

(SouthernBiotech, Birmingham, AL). Slides were stored at room temperature in the dark prior to imaging. Whole slice montage images were acquired with NIS-Elements imaging software on a Nikon Eclipse Ti or Ti2 Inverted Microscope System equipped with a motorized stage and epifluorescence illumination with standard DAPI, FITC, TRITC, Cy3, and Cy5 excitation/emission filter cubes. Confocal z stack images were acquired on an Olympus Fluoview 3000 laser scanning confocal microscope equipped with 488 nm, 543 nm, 594 nm, and 647 nm excitation laser lines.

Specificity of enhancer activity for the target striatal cell subclass or type was quantified as reporter and marker Ab double-positive neuron count divided by total reporter Ab positive neuron count and multiplied by 100 to obtain a percentage. The main ROI for analysis was in the center of the dorsal striatum region. A minimum of 50 neurons were counted in each ROI. Completeness of labeling of a given target striatal cell subclass or type was quantified as reporter and marker Ab double-positive neuron count divided by total marker positive neuron count and multiplied by 100 to obtain a percentage.

### RNAscope and IHC Co-Detection

#### Mouse

We performed RNA mFISH and IHC Co-Detection using the RNAscope Multiplex Fluorescent v2 and Co-Detection kits (Advanced Cell Diagnostics) with minor adjustments to the manufacturer’s protocols. . For the v2 kit, 30μm sections were prepared from mice perfused with 4% PFA, the brain extracted and sunk in 30% sucrose before OCT embedding. All coronal sections for both kits were cut using a cryostat (CM3050 S) and mounted on SuperFrost slides (ThermoFisher Scientific). Slides were stored at −80°C and used within one month.During sample preparation and pretreatment, we performed the hydrogen peroxide wash first before allowing the sections to dry for 2 h on the slide. The slide baking step was increased from 30 min to 1 h for improved tissue adherence. We added an additional 15-min baking step after the EtOH washes also for improved tissue adherence. Due to loss of native SYFP2 signal from the v2 kit protocol, we incorporated the RNA-Protein Co-detection kit to perform IHC to visualize the enhancer-driven SYFP2 while using the probes for the marker gene mRNA. We followed the co-detection kit protocol and placed the tissue in primary antibody solution at 4°C after the antigen retrieval step where it incubated overnight. The primary antibodies used where chicken anti GFP (1:250, Aves Lab Cat# GFP1020). The secondary antibody incubation occurs after the final HRP blocking step. The tissue was placed in secondary antibody solution for 3 h at room temperature. The secondary antibody used was goat anti chicken Alexa Fluor 488 (1:500, Invitrogen Cat#A11039.). The following probes were used: PVALB (ACD Cat#421931-C2) and PTHLH (ACD Bio Cat#456521-C3)To visualize the probes, we used the following TSA dyes and concentrations from ACD:, TSA Vivid 570 (1:1500, Cat#7535) for PVALB, and TSA Vivid 650 (1:1000, Cat#7536) for PTHLH. DAPI (Cat#323108) labeled nuclei. All slides are mounted with ProLong Gold Antifade mounting media (ThermoFisher Scientific Cat#P36930) and allowed to dry overnight before imaging on a Nikon Eclipse Ti2 Inverted Microscope System equipped with a motorized stage and epifluorescence illumination with standard DAPI, FITC, TRITC, Cy3, and Cy5 excitation/emission filter cubes. Specificity of enhancer-AAVs was calculated as: GFP+PVALB+PTHLH+/GFP+. Completeness was calculated as GFP+/PVALB+/PTHLH+/PVALB+/PTHLH+.

### SMART-seq v4 sample preparation and analysis (scRNA-seq)

Single-cell suspensions from enhancer-AAV RO injected mice were prepared for flow cytometry and single-cell RNA-seq from brain tissue as previously described. Briefly, for flow cytometry, we perfused mice transcardially under anesthesia with ACSF.1. We harvested the brains, embedded in 2% agarose in PBS, then sliced thick 350μm sections using a Compresstome (Precisionary Instruments) with blockface imaging, then picked the sections containing the dorsal striatum and dissected it out. We then treated dissected tissues with 30U/mL papain (Worthington LK003176) in ACSF.1 containing 30% trehalose (ACSF.1T) in a dry oven at 35°C for 30 min. After papain treatment we quenched digestion with ACSF.1T containing 0.2% BSA, triturated sequentially using fire-polished glass pipettes with 600, 300, and 150 micron bores, filtered the released cell suspensions into ACSF.1T containing 1% BSA, centrifuged cells at 100g for 10 min, then resuspended cells in ACSF.1T containing 0.2% BSA and 1 μg/mL DAPI prior to flow cytometry and sorting on a FACSAria III (Becton-Dickinson). Sample preparation for SMART-Seq was performed using the SMART-Seq v4 kit (Takara Cat#634894) as described previously. Single cells were sorted into 8-well strips containing SMART-Seq lysis buffer with RNase inhibitor (0.17 U/μL; Takara Cat# ST0764) and were immediately frozen on dry ice for storage at −80°C. SMART-Seq reagents were used for reverse transcription and cDNA amplification. Samples were tagmented and indexed using a NexteraXT DNA Library Preparation kit (Illumina Cat#FC-131-1096) with NexteraXT Index Kit V2 Set A (Illumina Cat#FC-131-2001) according to manufacturer’s instructions except for decreases in volumes of all reagents, including cDNA, to 0.4 x recommended volume. Full documentation for the scRNA-seq procedure is available in the ‘Documentation’ section of the Allen Institute data portal at http://celltypes.brain-map.org/

Samples were sequenced on an Illumina HiSeq 2500 as 50 bp paired end reads. Reads were aligned to GRCm38 (genecode v23) using STAR v2.7.1 with the parameter “twopassMode,” and exonic read counts were quantified using the GenomicRanges package for R as described previously (Tasic et al. 2018). To determine the corresponding cell type for each scRNA-seq dataset, we utilized MapMyCells hierarchical mapping from the Allen Institute https://portal.brain-map.org/atlases-and-data/bkp/mapmycells according to the posted instructions against the 10x Whole mouse brain taxonomy (CCN20230722). We applied quality control steps to exclude cells from the dataset if they had less than 1,000 genes detected and if they clearly did not map to BG cell subclasses corresponding to the dissected brain regions as described previously (Hunker et al. 2025).

### Macaque *in vivo* enhancer-AAV injections and histology

#### Surgeries

All procedures used with macaque monkeys conformed to the guidelines provided by the US National Institutes of Health and were approved by the University of Washington Animal Care and Use Committee. Three southern pig-tailed macaques (*Macaca nemestrina*) were used in this study for single *in vivo* injection of iodixanol gradient purified enhancer-AAV vectors using the Brainsight robotic surgery system (Rogue Research) (animal one, a 5-year-old 5 kg female, animal two, a 10-year-old 5kg male, animal three, a 3-year-old 5kg male). Brain MRIs for each animal were acquired and utilized for pre-planning of virus injection surgery using the Brainsight vet robot software v2.5 (Rogue Research). On the day of surgery, the animals were deeply anesthetized and positioned with their heads in the stereotaxic frame of the Brainsight surgery platform. A skin incision was made to expose the top of the skull, and burr hole craniotomies were drilled over the planned injection trajectory locations under the guidance of the Brainsight robotic surgery arm. After craniotomy, a prefabricated steel cannula with PE tubing attachment was loaded with a total of 20 μL AAV vector solution and attached to the robotic arm. For animal one, a total of 3μL of enhancer AiE0888m_C4 driving SYFP2 (AiP13863) AAV (3.00E+13 vg/mL) was injected into substantia nigra. For animal two, a total of 10µl of enhancer AiE1419m driving SYFP2 (AiP15043) AAV was injected into right caudate, and 5µl of enhancer AiE1223qrh driving SYFP2 (AiP14609) AAV was injected into right NAc/OT (both 2.00E+13 vg/mL). Animal three received a total of 5µl of enhancer AiE1426m driving SYFP2 (AiP15050) AAV (2.00E+13 vg/mL) into right caudate, and 3µl of enhancer AiE1485q driving SYFP2 (AiP15229) AAV was injected into right NAc/OT (1.00E+13 vg/mL). For each injection track, the cannula was advanced to the starting point of the injection target. Then, 1 μL of virus was gradually injected at each of five depths with 1mm spacing as the cannula tip was retracted dorsally. At the end of the final injection point in each track, the cannula was left in place for 10 min to allow virus diffusion prior to extracting the cannula from the brain. Note that additional virus injection sites were made in other distal brain locations in each macaque, but striatal labeling remained physically clearly separated from distal injection sites. At 35–40 days post-injection, each animal was sacrificed for brain collection and tissue expression analysis. The animals were transcardially perfused with 2L NMDG-aCSF solution. The brain was then removed and rapidly transported from the WaNPRC to the Allen Institute in chilled NMDG-aCSF solution for further tissue processing. The brain was first hemisected and drop-fixed in freshly prepared 4% PFA in PBS for 48 h at 4°C. After PFA fixation, the brain was transferred into 1XPBS +0.01% sodium azide solution. The brain hemispheres were examined and then 0.5 cm thick coronal slabs were cut through the striatum regions. Serial brain slabs were trimmed to fit into the wells of a six well tissue culture plate and submerged in 30% sucrose solution for a minimum of 48 h before further processing for histological analysis. Tissue slabs were mounted on the stage of a freezing-sliding microtome (Leica model SM2010R) on a bed of OCT and sub-sectioned to 30 μm thickness and stored in 1XPBS +0.01% sodium azide solution. Sections in the region of interest were selected for propidium iodide and DAPI staining and then slide mounted on 2”x3′′ glass slides. Alternatively, some tissue sections were used for free-floating immunostaining and RNAscope. Sections were dried onto slides on a slide warmer and cover-slipped with Vectashield Hardset mounting medium (Vector Labs) for PI and DAPI plus native SYFP2 imaging or Prolong Gold Antifade mounting medium (Life Technologies) for antibody staining experiments.

#### Immunohistochemistry

Brain slices were fixed in 4% PFA in phosphate buffered saline (PBS) at 4°C overnight or up to 48 h and then transferred to 1XPBS with 0.01% sodium azide as a preservative. Fixed slices were thoroughly washed with PBS to remove residual fixative, then blocked for 1 h at room temperature in 1XPBS containing 5% normal goat serum and 0.2% Triton X-100. After blocking, slices were incubated overnight at 4◦C in blocking buffer containing one or more of the following primary antibodies: chicken anti-GFP (Aves, 1:1000 or 1:2000), mouse anti-ChAT IgG2b for cholinergic cells (Atlas Labs, 1:1000), rabbit anti-nNos C-terminal for Sst-Chodl cells (Immunostar, 1:1000), and rabbit anti-TH for dopaminergic cells (Millipore, 1:1000). Following the overnight incubation, slices were washed for 15 min three times with 1XPBS and then incubated for 2 h at room temperature in dye-conjugated secondary antibodies including Alexa Fluor 488 (goat anti-chicken), Alexa Fluor 555 (goat anti-mouse), and Alexa Fluor 647 (goat anti-rabbit), and DAPI nuclear staining at 5µg/ml. Slices were then washed for 15 min three times with 1XPBS. The slices were then dried on glass micro-scope slides and mounted with Fluoromount-G. (SouthernBiotech, Birmingham, AL). Slides were stored at room temperature in the dark prior to imaging. Whole slice montage images were acquired with NIS-Elements imaging software on a Nikon Eclipse Ti or Ti2 Inverted Microscope System equipped with a motorized stage and epifluorescence illumination with standard DAPI, FITC, Cy3, and Cy5 excitation/emission filter cubes. Specificity of enhancer activity for the target cell subclass or type was quantified as reporter and marker Ab double-positive neuron count divided by total reporter Ab positive neuron count and multiplied by 100 to obtain a percentage. The main ROI for analysis was in the core of expression of the respective striatum region. A minimum of 50 neurons were counted in each ROI.

#### RNAscope Multiplex Fluorescent v2 & IHC Co-Detection

The RNAscope Multiplex Fluorescent v2 kit was used in tandem with the RNA-Protein Co-detection Ancillary kit (ACD) with a few modifications. The tissue was cut using a freezing sliding microtome and mounted on SuperFrost slides (ThermoFisher Scientific). Slides were stored at −80°C and used within one month. During sample preparation and pretreatment, we performed the hydrogen peroxide wash first before allowing the sections to dry for 2 h on the slide. The slide baking step was increased from 30 min to 1 h for improved tissue adherence. We added an additional 15-min baking step after the EtOH washes also for improved tissue adherence. Due to loss of native SYFP2 signal from the v2 kit protocol, we incorporated the RNA-Protein Co-detection kit to perform IHC to visualize the enhancer-driven SYFP2 while using the probes for the marker gene mRNA. We followed the co-detection kit protocol and placed the tissue in primary antibody solution at 4°C after the antigen retrieval step where it incubated overnight. The primary antibodies used where chicken anti GFP (1:250, Aves Lab Cat# GFP1020). The secondary antibody incubation occurs after the final HRP blocking step. The tissue was placed in secondary antibody solution for 3 h at room temperature. The secondary antibody used was goat anti chicken Alexa Fluor 488 (1:500, Invitrogen Cat#A11039).

The probes used were: DRD1 (Cat#549041-C2), DRD3 (Cat#1075641-C3), PTHLH (Cat#1588521-C3), PVALB (Cat#1075601-C2), , and RXFP1 (Cat#466741-C3). The RXFP1 probe is based on the Macaca fascicularis aurea species. The DRD1 and PVALB probes are based on the Macaca mulatta species. All probe target regions had a greater than 99.9% similarity rate when compared to the associated Macaca nemestrina regions using Nucleotide BLAST.

To visualize the probes, we used the following TSA dyes and concentrations from ACD: TSA Vivid 570(1:1500, Cat#323272) and TSA Vivid 650 (1:1500, Cat#323273)DAPI (Cat#323108) labeled nuclei. All slides are mounted with ProLong Gold Antifade mounting media (ThermoFisher Scientific Cat#P36930) and allowed to dry overnight before imaging on a Nikon Eclipse Ti2 Inverted Microscope System equipped with a motorized stage and epifluorescence illumination with standard DAPI, FITC, TRITC, Cy3, and Cy5 excitation/emission filter cubes. To quantify specificity, images were loaded into FIJI where brightness/contrast was manually adjusted to eliminate background signal in all channels. Cells were then marked using the Cell Counter plugin based on positivity for GFP, DRD1, DRD3, PTHLH, PVALB, or RXFP1. Specificity was calculated as GFP+DRD1+DRD3+/GFP+ for the D1-ICjs, GFP+DRD1+RXFP1+/GFP+ for the D1-NUDAP clusters, and GFP+PTHLH+PVALB+/GFP+ for the PVALB-PTHLH neurons. Completeness was calculated as GFP+PTHLH+PVALB+/PTHLH+PVALB+. Due to the non-uniform distribution of the D1-ICjs and D1-NUDAP clusters in a given section of ventral striatum based on their physical distribution, it was not possible to determine completeness. Specificity for the D1-ICjs and D1-NUDAP clusters was calculated in an ROI of the labeled neurons approximating 250 cells. We counted off target labeling in areas adjacent to the D1-ICjs and D1-NUDAP clusters, with 5% of DRD1+ MSNs co-labeled with GFP in each injection site.

#### TF motif analysis and sequence models

Refer to the companion study (Johansen et al. 2025) for details of DeepMacaqueBG model training. TF-MINDI (De Winter et al. 2026) is a Python tool that helps analyze DNA sequence patterns learned by deep learning models. It finds and groups recurring motifs from model attribution scores, links them to known transcription factor families, and provides visualizations to interpret regulatory signals in genomic data. We used TF-MINDI to perform motif-level analysis by replacing the internal motif/DBD database with group-specific patterns learned from global peak analysis using the DeepMacaqueBG model. The motifs were also annotated using JASPAR non-redundant vertebrate motifs based on Pearson correlation. The identified patterns represent seqlets that best match previously reported atlas patterns, annotated by Pearson correlation to JASPAR motifs. These patterns do not represent direct transcription factor binding motifs, but rather sequence features that are correlated with known motif profiles.

#### ATAC specificity

Accessibility signal across genomic peaks was quantified using bigWig tracks. Peaks were supplied as a BED file containing chromosome, start, and end coordinates along with metadata. Duplicate peaks with identical genomic coordinates were removed before processing.

Each bigWig file was used to extract continuous signal values. For every peak *p* and sample *s*, signal was summarized as the total of all non-missing per-base values across the interval:

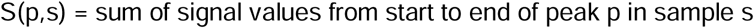

After computing raw signal values across samples, normalization was performed on a per-peak basis to calculate the relative contribution of each sample:

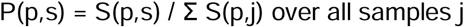

where P(p,s) represents the proportion of the total signal for peak *p* that originates from sample *s*. Peaks with zero total signal across all samples were retained and assigned zero proportions across samples.

The final output file contains the original peak metadata along with a proportion column for each BigWig sample. These normalized values can be used directly for downstream visualization (e.g., heatmaps, dot plots) and machine learning applications that require relative signal contributions per peak.

#### CREsted open chromatin prediction

Genomic enhancer regions were scored using DeepMacaqueBG CREsted convolutional neural network model. Peaks were supplied as a BED file containing chromosome, start, end, group, status, and peak ID columns.

A reference genome (Mmul10) was loaded using the DeepMacaqueBG CREsted Genome object, and the genome was registered for sequence retrieval. For each peak, the nucleotide sequence was fetched using the genomic coordinates. The DeepMacaqueBG CREsted model requires a fixed input length of 2,114 bp, so sequences shorter than this length were padded with “N” characters at the beginning and end to center the region within the model input window.

Each padded sequence was passed into the fine-tuned DeepMacaqueBG CREsted model to obtain prediction scores for all cell type classes simultaneously. The final file contains, for every genomic region, the original peak metadata (chromosome, start, end, group, status, peak ID) along with all DeepMacaqueBG CREsted model prediction scores.

#### PhyloP

Mean sequence conservation (phyloP) was computed for macaque ATAC-seq peaks by mapping orthologous coordinates to the human genome and extracting phyloP100way scores. Macaque peak regions were supplied as a BED file, and a separate BED file containing their corresponding human liftOver coordinates was used to pair macaque and human genomic intervals. Peaks were matched between the two files using the peak name field.

The human conservation signal was obtained from the phyloP100way BigWig file. For each peak, the script accessed the phyloP track and retrieved all per-base scores across the lifted-over human interval. Regions for which liftOver failed or for which coordinates were missing were assigned a conservation value of zero. To avoid artifacts due to erroneous liftOver expansion, any region with a lifted-over length greater than 5,000 bp was excluded from phyloP lookup and assigned a value of zero. For valid intervals, the mean conservation score was computed as the arithmetic mean of all non-missing phyloP values within the region.

#### Peak moments

To characterize the distribution of chromatin accessibility within each enhancer peak, we computed three statistical descriptors; skewness, kurtosis, and the bimodality coefficient across per-base ATAC-seq signal values for every cell type bigWig file.

First, candidate BG enhancer peaks were loaded from *peaks_NC_mapped_filtered.bed*. Peaks with duplicate genomic coordinates were removed. Missing or empty peak identifiers were replaced with unique IDs to ensure one identifier per genomic interval. TSS-normalized BigWig files representing ATAC-seq signal for each macaque BG cell type were retrieved from the project data directory.

For each peak, per-base accessibility values were extracted from every BigWig file using the pyBigWig Python library. Any missing values within a peak were replaced with zero. Peaks with completely uniform signal (all values identical or all zeros) were retained but assigned default statistics, because higher-order moments cannot be meaningfully computed for uniform distributions.

Skewness was used to describe asymmetry in the distribution of accessibility across the peak. Kurtosis quantified the overall peakedness or heaviness of the distribution. The bimodality coefficient was used to evaluate whether the signal profile within a peak tended toward unimodal or bimodal structure. All three metrics were computed using SciPy functions, and a custom implementation was used for the bimodality coefficient. Peaks with insufficient data points (fewer than four bases) were assigned missing values for this metric.

#### GC content calculation

We computed the GC content for each peak in the macaque dataset using the reference genome assembly (mmul10). Peaks were provided in BED format containing genomic coordinates and associated metadata. After reading the BED file, peak identifiers were standardized to ensure that every interval had a unique name. For each peak interval, we extracted the corresponding genomic sequence from the reference genome using pyfaidx, which enables efficient indexed retrieval of subsequences.

For each extracted sequence, we calculated the fraction of nucleotides that were either guanine or cytosine (GC content). The resulting GC content values were appended to the peak table and written to a tab-separated output file for downstream analysis.

