## Supplementary figures and images for "A Cross-Species Enhancer-AAV Toolkit for Cell Type-Specific Targeting Across the Basal Ganglia"

### Figure S1

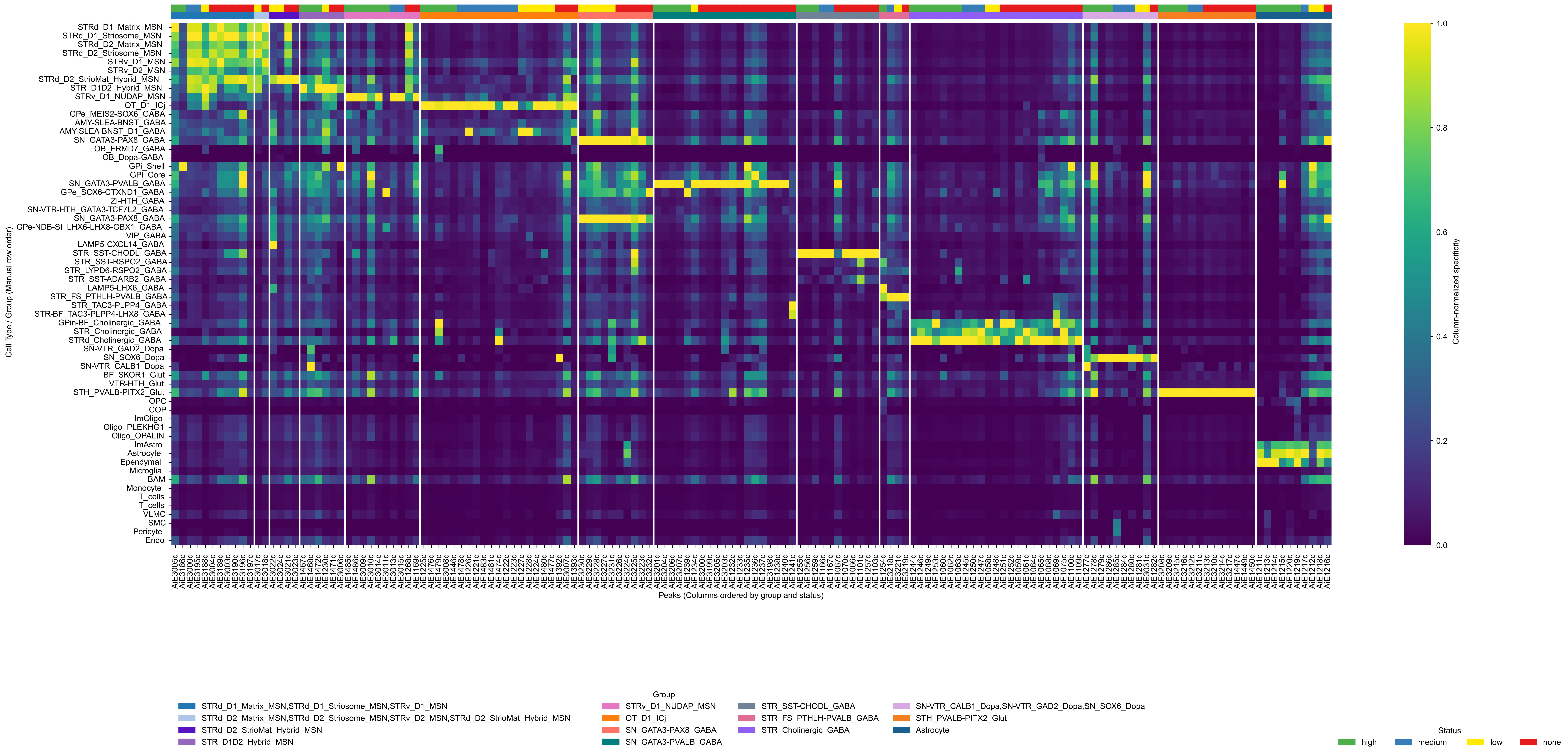

### Figure S2

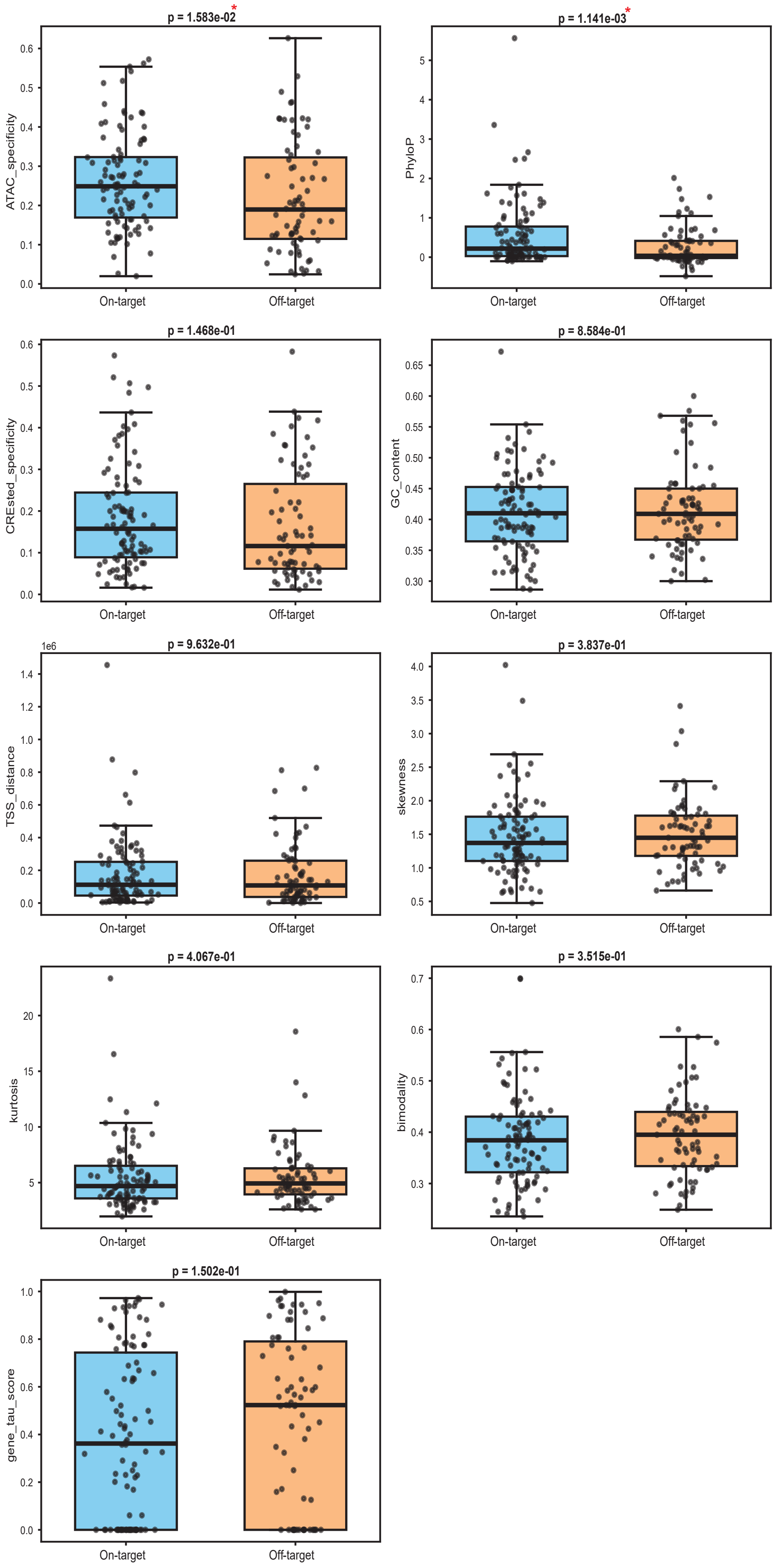

### Figure S3

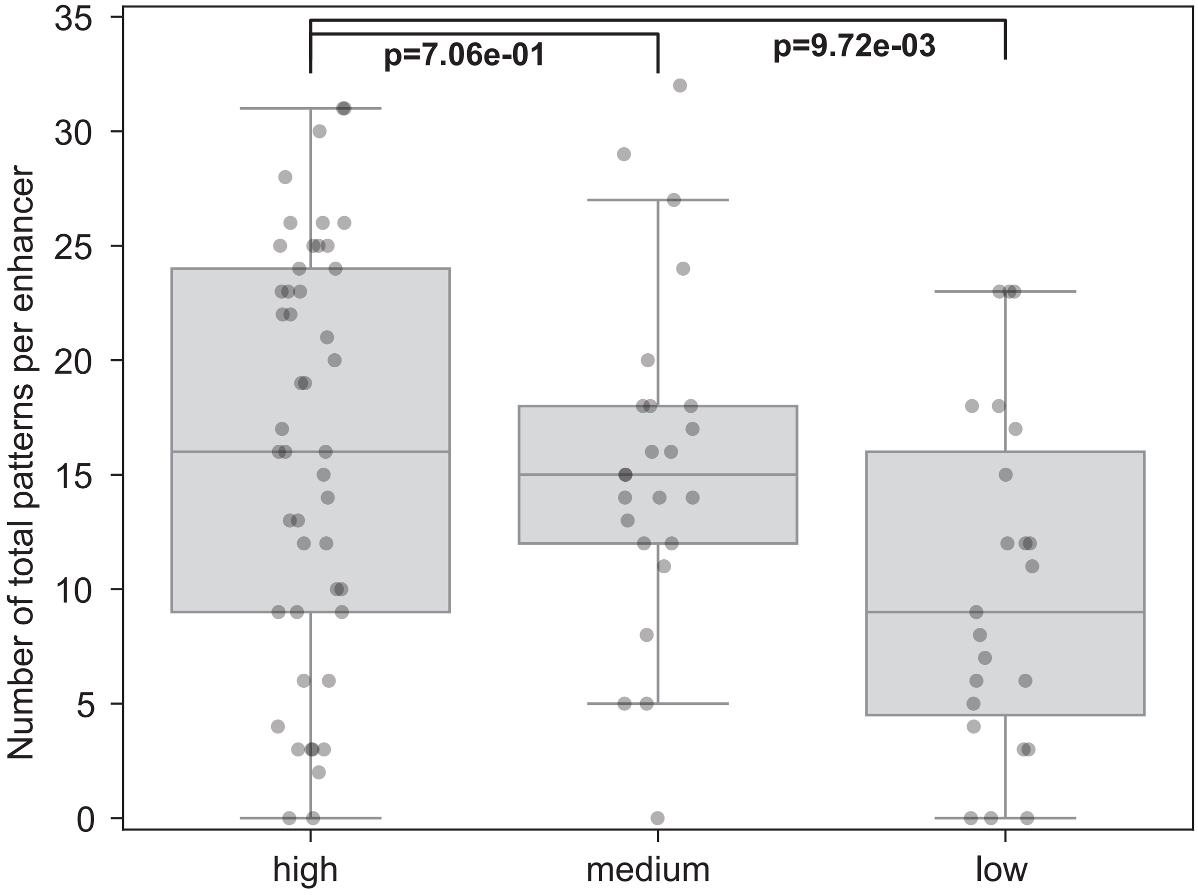
